# Prickle and Ror modulate Dishevelled-Vangl interaction to regulate non-canonical Wnt signaling during convergent extension

**DOI:** 10.1101/2023.08.29.555374

**Authors:** Hwa-seon Seo, Deli Yu, Ivan Popov, Jiahui Tao, Allyson Angermeier, Fei Yang, Sylvie Marchetto, Jean-Paul Borg, Bingdong Sha, Jeffrey D. Axelrod, Chenbei Chang, Jianbo Wang

## Abstract

Convergent extension (CE) is a fundamental morphogenetic process where oriented cell behaviors lead to polarized extension of diverse tissues. In vertebrates, regulation of CE requires both non-canonical Wnt, its co-receptor Ror, and several “core members” of the planar cell polarity (PCP) pathway. PCP was originally identified as a mechanism to coordinate the cellular polarity in the plane of static epithelium, where core proteins Frizzled (Fz)/ Dishevelled (Dvl) and Van Gogh-like (Vangl)/ Prickle (Pk) partition to opposing cell cortex. But how core PCP proteins interact with each other to mediate non-canonical Wnt/ Ror signaling during CE is not clear. We found previously that during CE, Vangl cell-autonomously recruits Dvl to the plasma membrane and keeps Dvl inactive. In this study, we show that non-canonical Wnt induces Dvl to transition from Vangl to Fz. Pk inhibits the transition, and functionally synergize with Vangl to suppress Dvl during CE. Conversely, Ror is required for the transition, and functionally antagonizes Vangl. Biochemically, Vangl interacts directly with both Ror and Dvl. Ror and Dvl do not bind directly, but can be cofractionated with Vangl. Collectively, we propose that Pk assists Vangl to function as an unconventional adaptor that brings Dvl and Ror into a complex to serves two functions: 1) simultaneously preventing both Dvl and Ror from ectopically activating non-canonical Wnt signaling; and 2) relaying Dvl to Fz for signaling activation upon non-canonical Wnt induced dimerization of Fz and Ror.

## Introduction

Throughout the animal kingdom, convergent extension (CE) is a universal morphogenetic engine that reshapes tissues during embryogenesis (Davey and Moens, 2017; Goodrich and Strutt, 2011; Huebner and Wallingford, 2018; Keller, 2002). Through polarized cell intercalation, directional cell migration or oriented cell division, CE generates powerful morphogenetic force to elongate a tissue in one direction while simultaneously narrowing it in the perpendicular direction. Disruption of CE can disturb normal embryogenesis from flies to mammals and causes various congenital disorders, including neural tube defects and skeletal disorders such as Robinow Syndrome and Brachydactyly type B (Butler and Wallingford, 2017; Wang et al., 2012; Yang and Mlodzik, 2015).

In vertebrates, CE is regulated by genes in the planar cell polarity (PCP) pathway. PCP refers to cell polarity orthogonal to that of apical-basal in epithelial cells. It was initially discovered in *Drosophila* as a signaling mechanism coordinating polarized cellular structures in the plane of the epithelium. The PCP pathway consists of six “core” proteins in flies, including three transmembrane proteins (the atypical cadherin Flamingo (Fmi), the receptor Frizzled (Fz), and the four-pass transmembrane protein Van gogh (Vang; Vangl in vertebrates)), and three cytoplasmic proteins (Dishevelled (Dsh; Dvl in mammals), Diego (Dgo), and Prickle (Pk)). A key feature of the core PCP proteins is that they assemble into two distinct complexes, those of Fmi/Fz/Dsh/Dgo and Fmi/Vang/Pk, that localize asymmetrically on opposing cell cortexes. Extensive genetic and imaging studies in flies, combined with computational modeling, have led to a model of feedback interaction in establishing core PCP protein distribution. The model proposes that Fmi on neighboring cells can establish homophilic interaction to facilitates cross talk between extracellular Fz and Vang in trans at the cell-cell junctions, with Dsh, Dgo and Pk function to stabilize the interacting complexes across the cell junctions. At the same time, these cytoplasmic proteins destabilize the juxtaposition of Fz and Vang in the same cell to segregate the complexes. Several mechanisms are used to facilitate both positive and negative feedback regulations, including selective interaction with partner proteins, post-translational modification of different components, stability of core proteins at cell junctions, transport of components along cytoskeleton, and membrane protein recycling and subcellular localization (Cho et al., 2015; Ressurreicao et al., 2018; Shimada et al., 2006; Strutt et al., 2013; Warrington et al., 2017). These feedback mechanisms act together to promote asymmetric clustering of Fz/Dsh/Dgo and Vang/Pk complexes at the distal and proximal cell junctions, respectively, to regulate asymmetric cytoskeletal organization and to coordinate planar polarity across the entire epithelium (Amonlirdviman et al., 2005; Axelrod and Tomlin, 2011; Humphries and Mlodzik, 2018; Strutt et al., 2016).

Though the PCP components are conserved in vertebrates, they coordinate cellular polarity not only in the plane of epithelial cells, but also in actively migrating cells, including neurons, neural crest, metastatic cancer cells, and cells undergoing CE. These cells share dynamic behaviors with constantly changing cell-cell contacts and interactions. Asymmetric localization of individual PCP proteins has been reported in a number of such cells, but the pattern varies and segregation of PCP complexes has not been consistently observed (reviewed in (Davey and Moens, 2017)). Moreover, as the majority of these studies focus on the activities of individual PCP component in regulating polarized cell behaviors, less is known about how PCP proteins interact with each other to coordinately control the migratory processes (Davey and Moens, 2017).

Potential differences in PCP protein interaction and function in *Drosophila* and vertebrates have emerged from some recent studies. Vangl1/2 were reported to function in a Celsr-independent manner to regulate mammalian airway morphogenesis (Paramore et al., 2024). On the other hand, whereas mutual inhibition between *Vangl* and *Fz/Dvl* is expected based on the *Drosophila* work, a number of reports also reveal a functional synergy between these proteins in mice. For instance, simultaneous decrease in gene dosage of *Vangl2* and *Dvl* enhanced CE defects in neural tube closure and cochlea elongation, and compound mouse mutants in *Vangl2* and several *Fz* genes show similar more exacerbated defects than those with mutations in individual genes (Etheridge et al., 2008; Wang et al., 2006; Yu et al., 2010; Yu et al., 2012). Upstream of Fz, the requirement of Wnt ligand in fly PCP signaling was debated initially (Chen et al., 2008; Wu et al., 2013), and disproved more recently (Ewen-Campen et al., 2020; Yu et al., 2020). In contrast, non-canonical Wnts, including Wnt5a and 11, are essential for CE in vertebrates (Grumolato et al., 2010; Heisenberg et al., 2000; Yamaguchi et al., 1999), and a functional synergy between Vangl2 and Wnt5a has been shown in development of multiple tissues in mice (Gao et al., 2011; Qian et al., 2007; Sinha et al., 2012; Wang et al., 2011). These findings, which collectively imply a positive role of Vangl in Wnt/Fz/Dvl mediated non-canonical Wnt signaling activation, bring up an essential question on how Vangl may both cooperate with and inhibit non-canonical Wnt/Fz/Dvl.

A further complication of non-canonical Wnt/ PCP signaling during vertebrate development is the involvement of several co-receptors including Ror1/2, Ptk7 and Ryk, whose functions are not linked to fly PCP ((Ripp et al., 2018); reviewed in (Green et al., 2014)). For instance, Ror2 has been shown to bind to Wnt5a together with Fz and is required to mediate Wnt5a-induced phosphorylation of Dvl in mammals (Grumolato et al., 2010; Ho et al., 2012; Nishita et al., 2010). Mouse mutants deficient in Ror1/2 phenocopy many defects of *Wnt5a* mutants (Ho et al., 2012), demonstrating a critical function of this co-receptor family in Wnt5a/ PCP signaling. Intriguingly, reduced gene dosages of both *Ror2* and *Vangl2* can lead to more severe morphogenesis defects than mutants of each individual genes, revealing a functional synergy between Vangl2 and Ror2 (Gao et al., 2011). This is reminiscent of the functional synergy observed between Vangl2 and non-canonical Wnt/Fz/Dvl. Ror2 was reported to interact with Vangl2 biochemically, and proposed to form a receptor complex with Vangl2 in response to Wnt5a (Gao et al., 2011). However, the biochemical and cell biological activities of the Ror2/ Vangl complex and how this may affect Wnt/Fz/Dvl PCP signaling is not understood in detail.

To understand feedback regulation of core PCP proteins in vertebrate CE, we have used the mouse and the *Xenopus* models to investigate functional and biochemical interactions of these proteins. Our previous work suggested that Vangl2 has dual activity in modulating Dvl function: it binds and recruits Dvl to the plasma membrane cell-autonomously and keeps it inactive, but at the same time enriching Dvl at this subcellular domain for Fz signaling upon stimulation by the Wnt11 ligand, which triggers release of Dvl from Vangl2 (Seo et al., 2017). In the current study, we attempted to address two questions raised by this model: 1) how will Vangl’s molecular partner Pk modulate Vangl-Dvl interaction during CE; and 2) if Dvl is sequestered by Vangl, how can it gain access to Fz in the response to Wnt?

In flies, Pk clusters with Vang to the proximal cell junction and is required to generate feedback amplification for asymmetric localization of both Vang and Dsh/Fz. Pk is shown to stabilize Fz-clusters on the plasma membrane in neighboring cells, but destabilizes Fz-clusters in a Dsh-dependent manner through endocytosis in the same cell (Warrington et al., 2017). Competitive binding of Pk to Dsh to prevent its plasma membrane recruitment by Fz has been proposed as a mechanism to destabilize Fz/Dsh clustering in the same cell (Tree et al., 2002). But binding between Pk and Dsh was reported to be quite weak (Bastock et al., 2003), and over-expressing Pk in *Xenopus* cannot effectively abolish Fz7-mediated recruitment of Dvl to the plasma membrane (Takeuchi et al., 2003; Veeman et al., 2003). These studies thus do not fully support the notion of competitive binding to Dvl as the underlying mechanism for Pk to destabilize Fz clusters or its action during CE.

In this study, we used gastrulating *Xenopus* embryos as a CE model to carry out functional, biochemical and imaging studies, and found that Pk helps Vangl2 to sequester both Dvl2 and Ror2, whereas Ror2 is needed for Dvl to transition from Vangl to Fz in response to non-canonical Wnt. We propose a novel model in which Pk assists Vangl to function as an unconventional adaptor that brings Dvl and Ror2 into a complex to serves two functions: 1) simultaneously preventing both Dvl and Ror2 from ectopic activation; and 2) relaying Dvl to Fz via Ror2 upon non-canonical Wnt activation. We propose that these two actions together help to modulate the threshold and dynamics of signaling activation in response to non-canonical Wnt.

## Results

### Pk synergizes with Vangl2 to suppress Dvl during CE

To probe the functional network of core PCP proteins during CE, we first studied how Pk interacts with Vangl2 and Dvl to regulate *Xenopus* body axis elongation in both gain- and loss-of-function scenarios. To overexpress Pk in *Xenopus* embryos, we used two different mRNAs that encode either a GFP tagged mouse Pk2 (mPk2) or a Flag-tagged *Xenopus* Pk1 (XPk) (Takeuchi et al., 2003; Vladar et al., 2012). Similar to the previous report (Takeuchi et al., 2003), we found that injecting *Xpk* or *mPk2* mRNA into the dorsal marginal zone (DMZ) of 4-cell stage *Xenopus* embryos can block CE in a dose dependent manner (Suppl. Fig. 1a). Quantification of the length-to-width ratio (LWR) indicates that 1 and 2 ng *pk* mRNA injection can reproducibly cause moderate and severe CE defects, respectively, but 0.5 ng *Xpk* only results in a slight LWR reduction that is not statistically significant (Suppl. Fig. 1a’).

With co-injection of mRNAs, however, even a very small dose of 0.1 ng *Xpk* is sufficient to cause sever CE defects together with 0.1 ng mouse *Vangl2* (*mVangl2*), which produces only a moderate CE defect when injected by itself (Fig. 1a, a’). Similarly, co-injecting 0.25 ng *mPk2*, which causes no CE defects by itself, also significantly enhances the CE defect induced by 0.1 ng *mVangl2* (Fig. 1b, b’).

**Figure 1.**
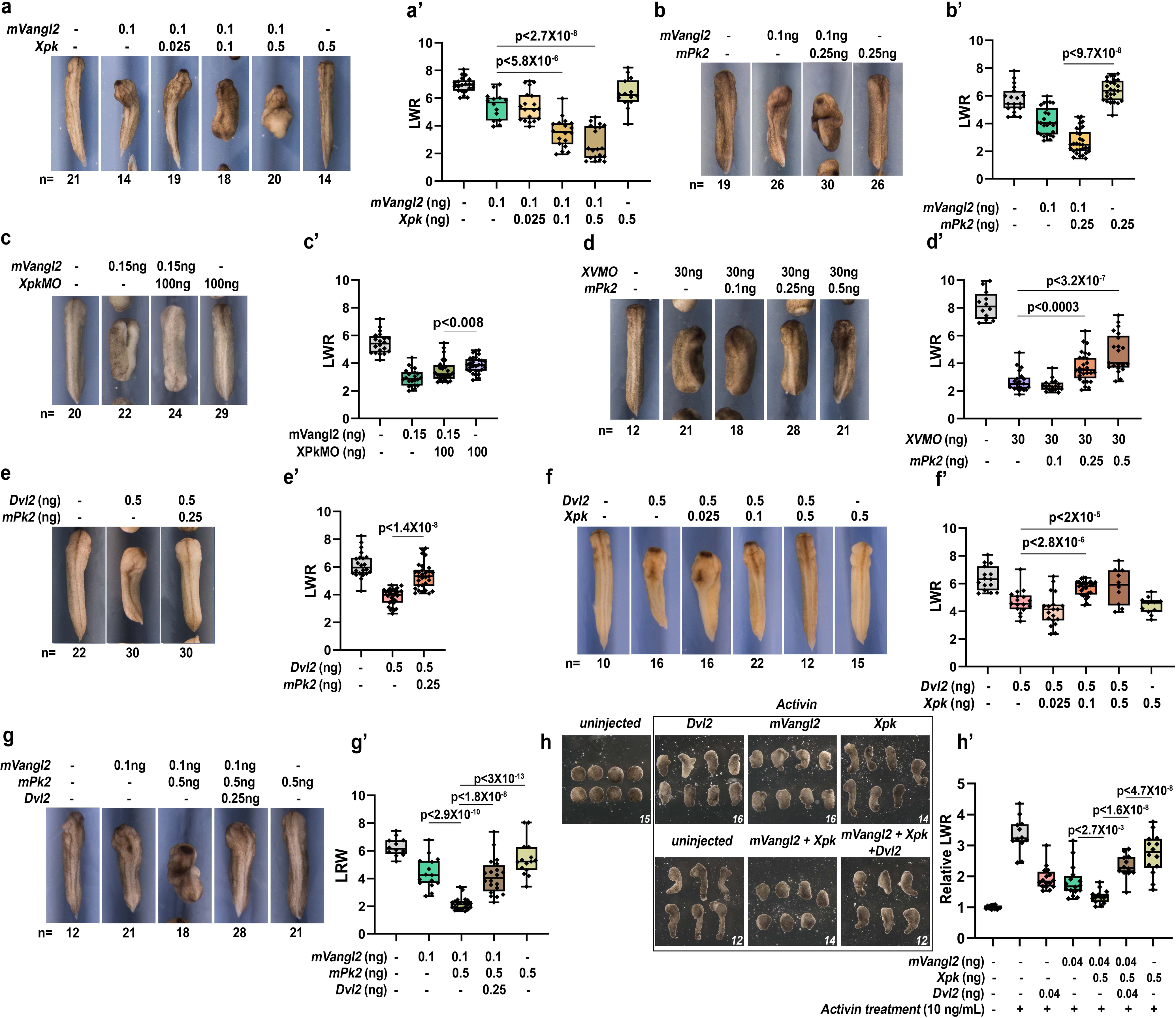
Pk synergize with Vangl to suppress Dvl during CE. Injecting 0.1 ng mouse *Vangl2* mRNA (*mVangl2*) into the DMZ results in moderate CE defects and reduction of the length-to-width ration (LWR), and the phenotypes are significantly enhanced by co-injecting a small dose of *Xpk* (a, a’) or *mPk2* (b, b’) mRNA that causes minimal or no CE defect *per se*. On the other hand, higher dose *mVangl2* (0.15 ng) induced more severe CE defects can be rescued by knocking down endogenous Pk using *Xpk*MO (c, c’); whereas knockdown of endogenous *vangl2* (*XV*MO) induced CE defects can be rescued by moderate mPk2 overexpression (d, d’). Conversely, 0.5 ng mouse *Dvl2* injection induced CE defects can be dose-dependently rescued by *mPk2* (e, e’) or *Xpk* (f, f’) co-overexpression; and mVangl2-mPk2 co-overexpression induced severe CE defect can be rescued by co-injecting Dvl2 (g, g’). Similarly, in activin-induced animal cap elongation assay, mVangl2 synergizes with Xpk to induce strong CE defect, which can be rescued significantly by co-injection of mouse Dvl2 (h, h’). CE phenotype was determined by quantifying the length-to-width ratio (LWR) of the embryos or animal cap explants in each group (a, b, c, d, e, f, g, h). Experiments were repeated three times and the total number of embryos or explants analyzed is indicated below each panel in (a)-(h). Data are presented as box plots in (a’), (b’), (c’), (d’), (e’), (f’), (g’) and (h’), with the whiskers indicating the minima and maxima, the center lines representing median, the box upper and lower bounds representing 75th and 25th percentile, respectively. Two-tailed, unpaired T-test was used to compare the LWR of different groups, and the p vales are indicated in (a’)-(h’) between different groups.

Conversely, we found that knocking down endogenous XPk level using antisense morpholino (*Xpk*MO, Suppl. Fig. 1b-d’) can rescue *mVangl2* over-expression induced CE defect (Fig. 1c, c’), whereas over-expressing Pk can dose-dependently rescue *Xvangl2* morpholino (*XV*MO) knockdown-induced CE defect (Fig. 1d, d’). These gain- and loss-of-function results together demonstrate that Pk functionally synergize with Vangl during CE.

We then tested how Pk may affect Dvl function during CE. When co-injected, *mPk2* or *Xpk* can dose-dependently rescued the CE defects induced by *Dvl2* over-expression (Fig. 1e-f’), suggesting that Pk functionally antagonizes Dvl during CE. Together with our previous finding that Vangl exerts bimodal regulation of Dvl (Seo et al., 2017), these results suggest that Pk may synergize with Vangl to suppress Dvl function during CE. Consistent with this idea, we found that the severe CE defects induced by *Pk* and *Vangl2* co-injection could be rescued by overexpressing *Dvl2* (Fig. 1g, g’). Using activin-induced animal cap elongation as an additional assay for CE, we further confirmed that *Xpk* synergizes with *mVangl2* to induce severe CE defect, which can be rescued by co-overexpression of Dvl2 (Fig. 1h, h;).

### Vangl interaction with and recruitment of Pk to the plasma membrane is essential for their functional synergy

To understand how Pk synergizes with Vangl to represses Dvl, we first investigated whether they could modulate each other’s protein levels. It was reported that Vang could control Pk stability indirectly through ubiquitination in flies (Cho et al., 2015; Strutt et al., 2013), while in zebrafish Pk could down-regulate Dsh/Dvl protein level (Carreira-Barbosa et al., 2003). In *Xenopus* embryos or explants undergoing CE, however, we found that morpholino knockdown of *Xvangl2* or over-expression of *mVangl2* did not affect the protein level of co-injected mPk2 or XPk (Suppl. Fig. 2a, b, c). In contrast, co-transfecting XPk with Vangl2 in cultured HEK293T cells did lead to significant reduction of XPk protein level (Suppl. Fig. 2d, also see Pk2 down-regulation by Vangl2 in 293 cells in (Nagaoka et al., 2019)). Therefore, Vang/Vangl2-modulation of Pk stability seems to be context-dependent, and does not account for the observed synergy between Vangl2 and Pk during *Xenopus* CE. Furthermore, over-expression of Pk does not alter the protein level of co-injected Vangl2 or Dvl2 (Suppl. Fig. 2a, b), indicating that during *Xenopus* CE Pk does not synergize with Vangl2 or antagonize Dvl2 by altering their protein levels.

To explore other mechanisms that may explain how Pk synergizes with Vangl2 to antagonize Dvl during CE, we examined the effect of Vangl2 on Pk’s sub-cellular localization. When EGFP tagged mPk2 is expressed in either the animal cap or DMZ, it displays diffused cytoplasmic distribution and variable enrichment at the plasma membrane (Suppl. Fig. 3a, c; Suppl. Fig. 4a). Co-injection of mVangl2 significantly increased plasma membrane enrichment of mPk2 in both animal cap and DMZ explants (Suppl. Fig. 3b; Suppl. Fig. 4b, e). On the other hand, morpholino knockdown of endogenous *Xvangl2* diminished mPk2 plasma membrane enrichment, which could be restored by co-injection of a small amount of mVangl2 (Suppl. Fig. 3d, e; Suppl. Fig. 4c, d and e). Together, these data indicate that Vangl2 is both necessary and sufficient to recruit Pk to the plasma membrane.

To test whether plasma membrane recruitment of Pk by Vangl is important for their functional synergy, we took advantage of a Vangl2 R177H variant identified in a patient with diastematomyelia (Kibar et al., 2011). This variant changes the highly conserved Arg177 to a histidine in the intracellular loop region between TM2 (transmembrane domain) and TM3 (Fig. 2a). Importantly, this variant does not perturb Vangl2 plasma membrane trafficking/ localization (Fig. 2b, b’), its protein level (Fig. 2g, Suppl. Fig. 5e), or its ability to interact with and recruit Dvl to the plasma membrane (Suppl. Fig. 5a-d). The variant, however, reduced Vangl2 interaction with Pk and recruitment of Pk to the plasma membrane (Fig. 2c-g).

**Figure 2.**
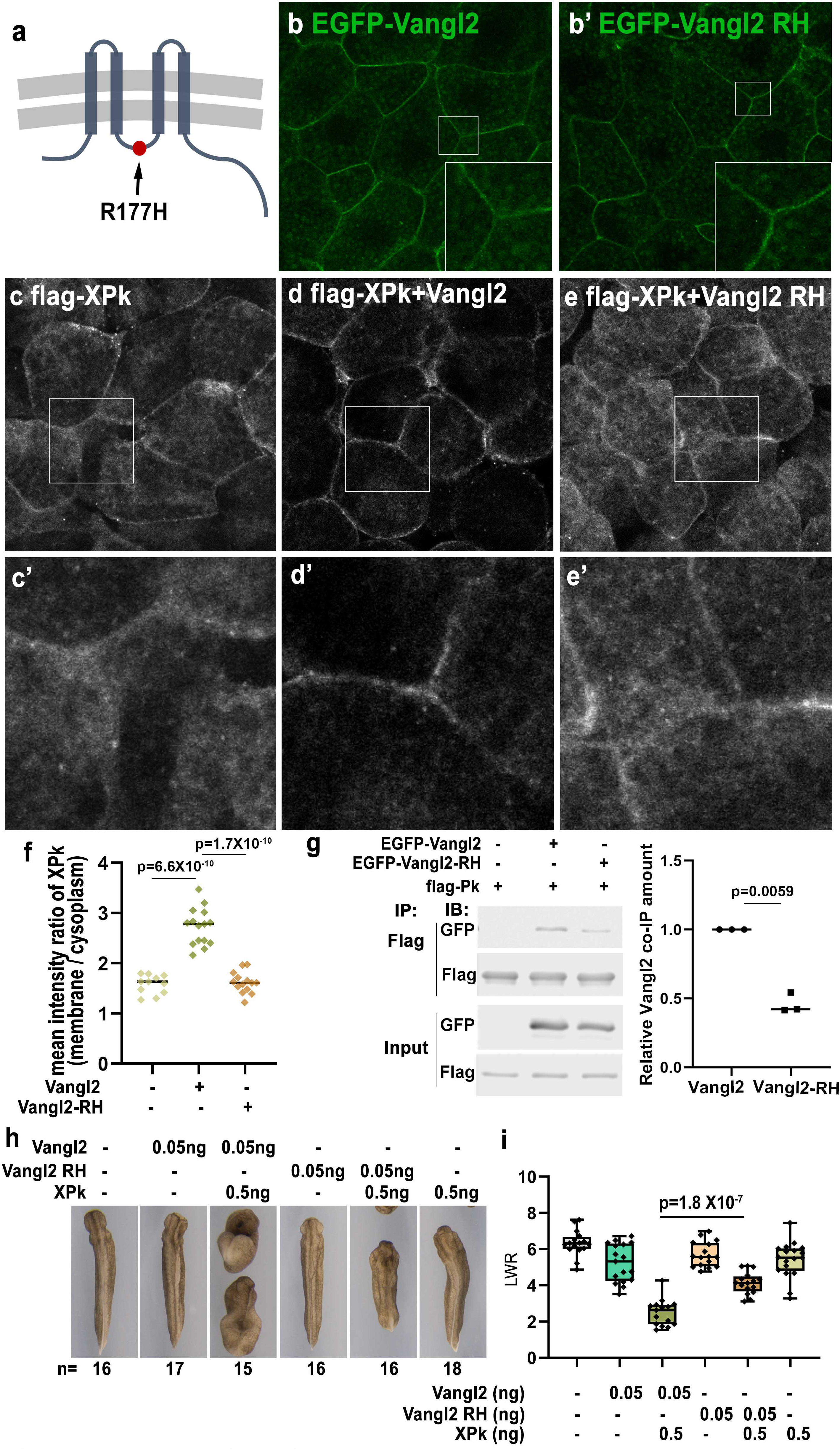
Vangl2 RH variant diminishes membrane recruitment of Pk and reduces functional synergy with Pk. Schematic illustration showing the structure of Vangl2 and location of the R177H variant at the intracellular loop region between transmembrane domain 2 and 3 (a). When expressed in *Xenopus* animal cap cells, EGFP-Vangl2 RH displays plasma membrane localization indistinguishable from wild-type EGFP-Vangl2 (b, b’). Immunostaining shows that flag-XPk displays diffuse cytoplasmic localization with some membrane enrichment when expressed alone (c, c’). The plasma membrane localization of flag-XPk is enhanced significantly by co-expression of wild-type Vangl2 (d, d’), but only modestly by Vangl2 RH variant (e, e’). (f) Quantification of the ratio of plasma membrane vs. cytoplasmic flag-XPk signal intensity in (c), (d) and (e). Co-IP and western blot show that R177H mutation does not alter Vangl2 protein level, but reduces binding to flag-XPk (g, n=3 biological repeats). Functionally, co-injection of 0.05 ng *Vangl2* and 0.5 ng *Xpk* can strongly synergize to disrupt CE, but the synergy is significantly reduced when *Vangl2 RH* variant mRNA is co-injected with *Xpk* (h). The CE phenotype was determined by quantifying the length-to-width ratio (LWR) of the embryos in each group in (h). Experiments were repeated three times and the total number of embryos analyzed is indicated below each panel in (h). Data are presented as box plots in (i), with the whiskers indicating the minima and maxima, the center lines representing median, the box upper and lower bounds representing 75th and 25th percentile, respectively. Two-tailed, unpaired T-test was used to compare the LWR of different groups, and the p vales are indicated between different groups.

Functionally, the R177H variant significantly reduced the synergy between Vangl2 and Pk during CE (Fig. 2h, i). Moreover, compared to wild-type Vangl2, Vangl2 R177H results in significantly less severe CE defects when over-expressed alone in the DMZ (Suppl. Fig. 6a, a’), and is less capable of suppressing Dvl-Fz mediated signaling activation during CE when co-expressed (Suppl. Fig. 6b, b’). We interpret these data to suggest that: 1) direct binding of Vangl2 to Pk, through which Pk is recruited to the plasma membrane, is crucial for their functional synergy during CE; and 2) direct binding of Vangl to Dvl alone may not be sufficient to suppress Dvl, and simultaneous interaction with Pk may be required for Vangl to efficiently inhibit Dvl during CE.

### Pk synergizes with Vangl to sequester Dvl from Fz

To investigate how Pk may help Vangl to suppress Dvl during CE, we first tested whether Pk over-expression may disrupt Dvl interaction with Fz. In both animal cap and DMZ cells, Fz7 can recruit mCherry tagged Dvl2 (Dvl2-mCh) to the plasma membrane (Suppl. Fig. 7b, e). Co-expression of GFP-mPk2, however, does not perturb Fz7-mediated plasma membrane recruitment of Dvl2 (Suppl. Fig. 7c, f), suggesting that over-expression of Pk alone cannot effectively disrupt Dvl-Fz interaction in *Xenopus*.

We then investigated the alternative possibility that Pk may help Vangl to sequester Dvl, and thereby preventing Wnt-Fz-Dvl interaction to inhibit non-canonical Wnt signaling activation during CE. Our previous studies provided evidence that Vangl recruits Dvl into an inactive complex at the plasma membrane; and Wnt11 can induce dissociation of Dvl from Vangl (Angermeier et al., 2025; Seo et al., 2017). We therefore first tested whether Pk may reinforce Vangl-Dvl interaction to counter the dissociation effect by Wnt11. When co-injected into the DMZ, mPk2 significantly prevented Wnt11-induced dissociation of Flag-Dvl2 from EGFP-Vangl2, although in the absence of Wnt11 co-injection it did not substantially increase Dvl2-Vangl2 interaction (Suppl. Fig. 8). These data suggest that, at least under the condition of our co-IP experiment, Pk may not directly impact the steady-state binding between Vangl and Dvl, but may strengthen Dvl sequestration by Vangl to inhibit its response to non-canonical Wnt ligand.

To investigate how Pk may help Vangl to regulate Dvl’s response to non-canonical Wnt, we performed imaging studies. In *Xenopus* and zebrafish, Wnt11 can induce formation of Fz-Dvl complexes that cluster as patches at the cell-cell contacts (Angermeier et al., 2025; Witzel et al., 2006; Yamanaka and Nishida, 2007). We therefore tested how Vangl/ Pk may affect formation of Wnt11-induced Fz-Dvl patches. In DMZ (Suppl. Fig. 9a-c’) or animal cap (Fig. 3a-c’) explants, co-injection of *Xenopus Wnt11* can induce Dvl2-EGFP, Dvl2-mScarletI (Dvl2-mSc) or Dvl2-mCh to form distinct patches along cell-cell contacts. By separate injection of *Dvl2-mSc* and *Dvl2-EGFP* into two adjacent blastomeres, we found that the Wnt11-induced Dvl2-mSc and Dvl2-EGFP patches are aligned along the adjacent cell borders (Suppl. Fig. 9a-c’). This result is consistent with the previous reports (Witzel et al., 2006; Yamanaka and Nishida, 2007), and indicates that Wnt11 induces symmetric clustering of Dvl across the border of adjacent cells, a scenario that is different from the asymmetric clustering of Fz/Dvl and Vang/Pk complexes between adjacent cells commonly observed during PCP establishment in epithelial tissues.

**Figure 3.**
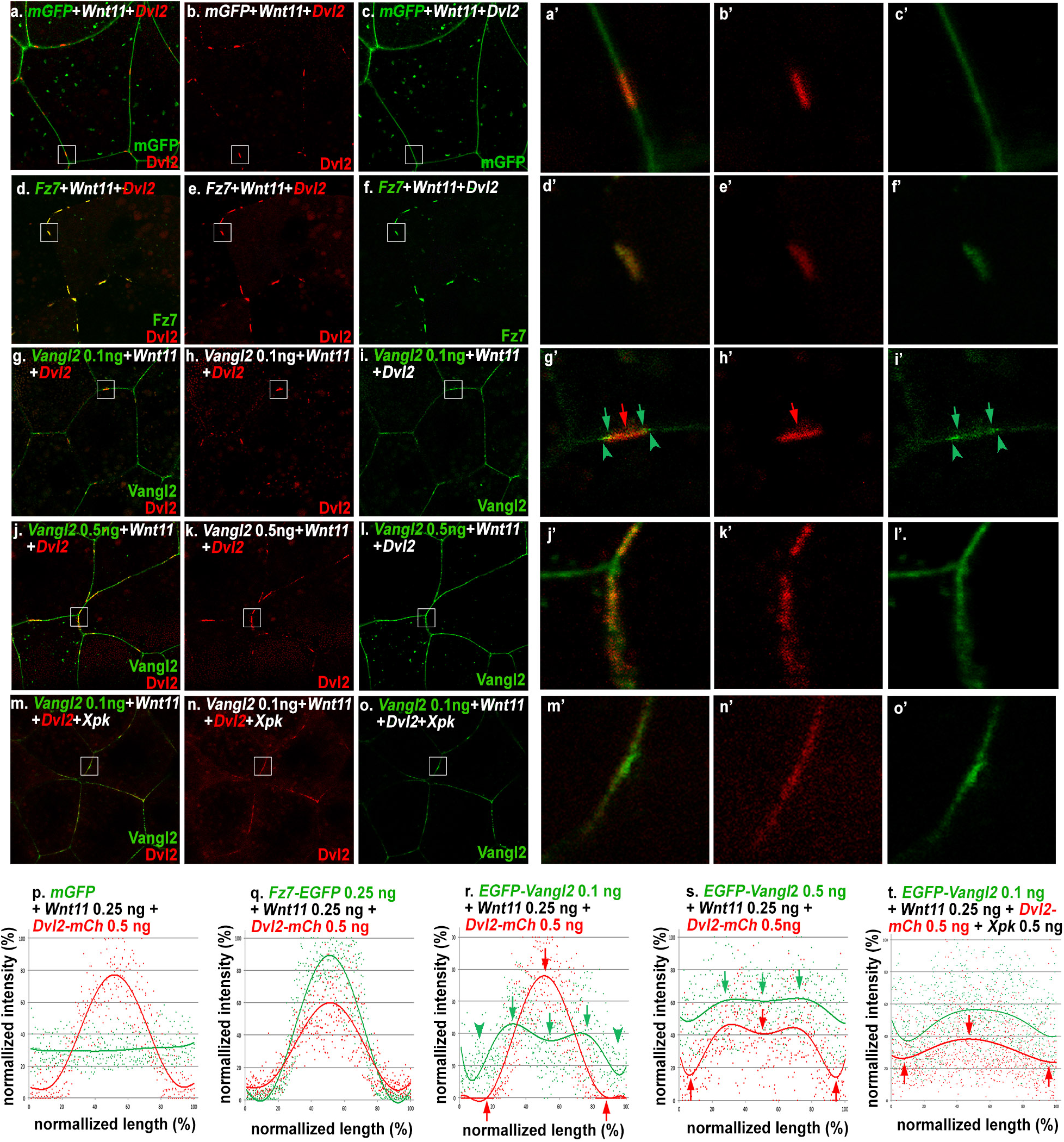
Pk synergizes with Vangl2 to inhibit Wnt11-induced formation of Dvl patches. In animal cap explants, 0.25 ng *Wnt11* injection with 0.5 ng of mCh-tagged mouse *Dvl2* (*Dvl2-mCh*) and membrane-GFP (*mGFP*) induces formation of distinct Dvl2 patches at the cell-cell contact (a-c’). Co-injection indicates that these Dvl2 patches completely overlap with Fz7-EGFP (d-f’). In contrast, Vangl2, when expressed at moderate levels (0.1 ng), is distributed more broadly along the plasma membrane (g-i), but also displays enrichment immediately outside and at the edge of Dvl2 patches (g’-I’, arrowheads and arrows, respectively). High level of *Vangl2* injection (0.5 ng) inhibits Wnt11-induced Dvl2 patch formation and makes Dvl2 more evenlydistributed with Vangl2 (j-l’). The same effect can also be achieved by co-expressing Pk with moderate level of Vangl2 (m-o’). (p-t) Measurement of the relative intensity of Dvl2 along the patches with either membrane GFP, Fz7, Vangl2 at moderate (0.1 ng) and high (0.5 ng) levels, or Vangl2 (0.1 ng) with *Xpk* (0.5 ng) co-injection.

When EGFP-tagged *Xenopus* Fz7 is co-injected in animal cap or DMZ explants, it exclusively forms membrane patches that completely overlapped with Dvl2 (Fig. 3d-f’; Suppl. Fig. 9d-f’), consistent with clustering of Fz with Dvl. Co-injecting moderate level of EGFP-tagged mouse Vangl2 (0.1 ng) does not perturb Wnt11-induced Dvl2 patch formation. In contrast to Fz7, Vangl2 is distributed broadly on the plasma membrane, but to our surprise it also displays overlapping enrichment with Dvl2 patches (Fig. 3g-i; Suppl. Fig. 9g-i). Close examination revealed that in most cases Vangl2 is enriched at the edges of Dvl2 patches, but diminished at the center (Fig. 3g’-i’ and Suppl. Fig. 9g’-i’, compare red arrows and arrowheads to green arrows and arrowheads). In 3-D reconstructed confocal images, enriched Vangl2 often forms rings that encircle Dvl2 patches (Suppl. Fig. 10c, white arrows; and enlarged views in a’-c’).

To quantitatively assess their spatial distribution, we measured and plotted the relative intensity of Dvl2 against Fz7, Vangl2 or membrane-GFP control along the length of ten representative patches in animal cap explants (Fig. 3p-r). Our analyses revealed that membrane-GFP displayed no enrichment along Wnt11-induced Dvl2 patches (Fig. 3p). Fz7, however, showed enrichment that correlated strongly with Dvl2: their intensities follow a similar pattern of increasing sharply from the edge and peaking coincidentally at the center of the patches (Fig. 3q). In contrast, Vangl2 enrichment starts to appear slightly outside of Dvl2 patches (Fig. 3g’, i’, r, green arrowheads), peaks at the edges as Dvl2 intensity begins to rise, and dips at the center where Dvl2 and Fz7 intensities reach the maximum (Fig. 3g’, i’, r, green arrows). Analyses of DMZ explants showed the same results (Suppl. Fig. 9q, r). These imaging analyses are consistent with our co-IP data (Suppl. Fig. 8), and further suggest the possibility that Dvl may leave Vangl and transition to Fz upon Wnt11 induction.

We then tested whether addition of Pk can help Vangl2 to counter the effect of Wnt11. Indeed, co-injecting Pk with 0.1 ng Vangl2 effectively reduced Wnt11-induced Dvl2 patch formation, making Dvl2 more evenly distributed along the plasma membrane (Fig. 3m-o; Suppl. Fig. 9m-o) and overlap with Vangl2 (Fig. 3m’-o’, t; Suppl. Fig. 9m’-o’, t). Similar reduction of Dvl patch formation can also be achieved, albeit less effectively, by over-expression of high level Vangl2 alone (Fig. 3j-l’, s; Suppl. Fig. 9j-l’, s).

To examine how Vangl2 and Pk could affect Fz7 enrichment in the Wnt11-induced Fz/Dvl patches, we co-injected fluorescent protein-tagged Dvl2 and Fz7 in both animal cap and DMZ explants (Fig. 4; Suppl. Fig. 11). In both cases, moderate over-expression of Vangl2 or Pk individually does not affect enrichment of Fz7 within Wnt11-induced Dvl2 patches (compare Fig. 4a-c’ with d-i’; Suppl. Fig. 11a-c’ with d-I’), but Vangl2 and Pk together not only perturb Dvl2 patches, but also disperse Fz7 into small puncta (Fig. 4j-l’; Suppl. Fig. 11j-l’). Close examination revealed that some of the Fz7 puncta are on the plasma membrane and largely co-localize with Dvl2. The rest of Fz7 puncta, however, are located in the cytoplasm near the plasma membrane and appear to be endocytosed vesicles (arrows in Fig. 4k’, l’ and Suppl. Fig. 11k’ and l’). Interestingly, these cytoplasmic puncta contain only Fz7 but not Dvl2 (compare arrows in Fig. 4j’ to k’; and arrows in Suppl. Fig. 11j’ to k’).

**Figure 4.**
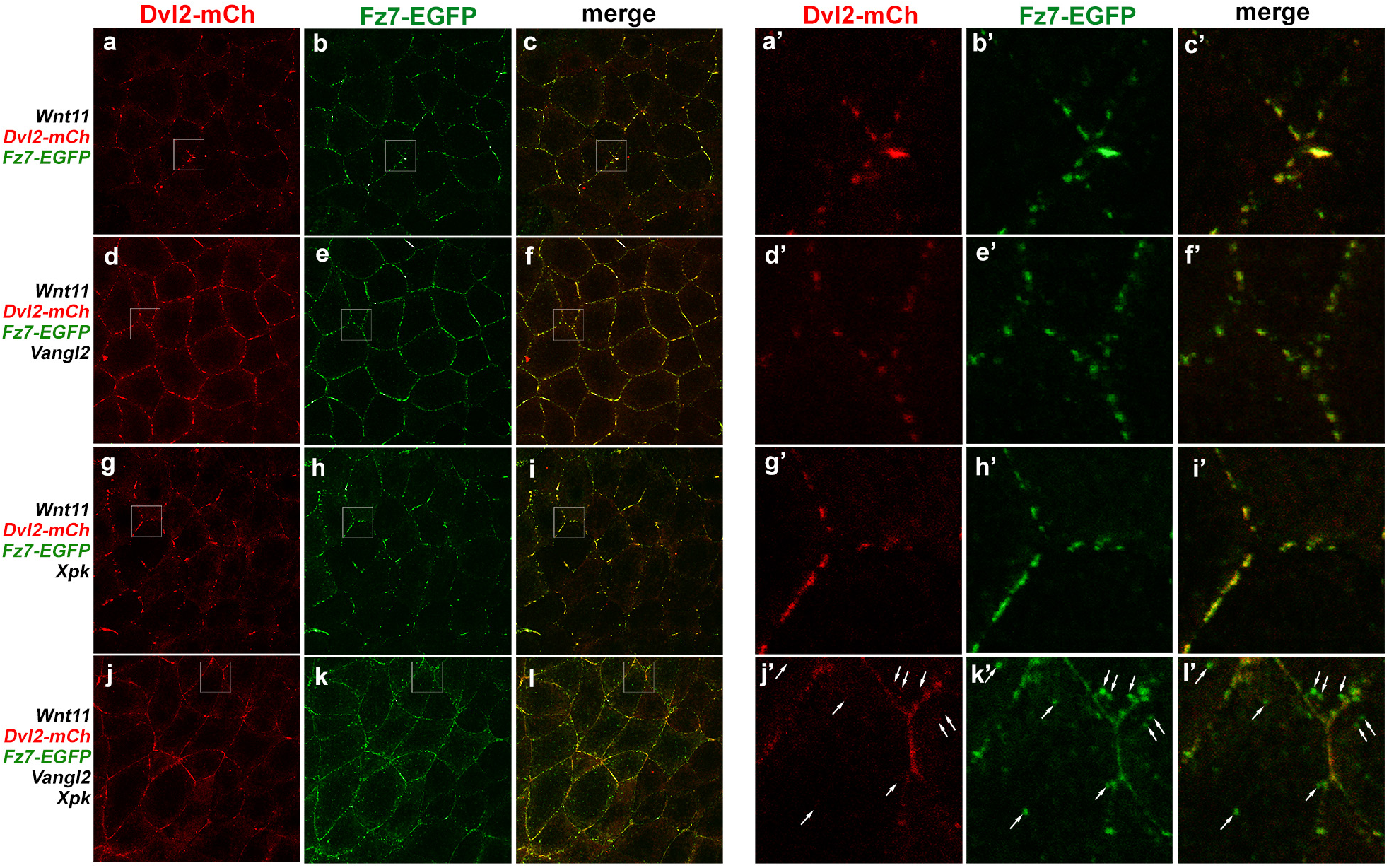
Pk helps Vangl2 to inhibit Wnt11-induced clustering of Fz7-Dvl2 complexes. In animal cap explants, Wnt11 induces formation of overlapping Fz7-EGFP and Dvl2-mCh patches at the cell-cell contact (a-c’). These patches are not affected by over-expressing moderate level of Vangl2 (d-f’, 0.1 ng mRNA) or XPk (g-i’, 0.5 ng mRNA) individually. Vangl2 and XPk co-expression, however, not only disrupt Dvl2-mCh patches but also disperse Fz7- EGFP patches into small puncta (j-l). Enlarged views revealed that some of the Fz7-EGFP puncta are on the plasma membrane and remain co-localized with Dvl2-mCh, while the others are located in the cytoplasm near the plasma membrane (k’’, l’, arrows) and contain only Fz7 but not Dvl2 (compare arrows in j’ to k’).

To confirm that the cytoplasmic Fz7 puncta are endocytosed vesicles, we performed FM4-64 dye uptake experiment (Cho et al., 2015; Classen et al., 2005). FM4-64 is a membrane impermeable fluorescent dye that can only be internalized through endocytosis. When we incubated the explants with FM4-64, we found that many Fz7 puncta induced by Vangl2/Pk co-injection were also positive for FM4-64 (Suppl. Fig. 12d-f’), indicating that they are indeed endocytic vesicles.

These data imply that Pk may assist Vangl to sequester Dvl, thereby reducing the accessibility of Dvl to attenuate Fz-Dvl complex formation in response to Wnt11, and resulting in Fz destabilization at the plasma membrane. To test this idea, we reduced Dvl availability at the plasma membrane using another strategy. Over-expression of DshMA, a mitochondrial tethered Dvl, can sequester endogenous Dvl to the mitochondria (and away from the plasma membrane) through DIX-domain mediated oligomerization (Park et al., 2005). We found that DshMA injection indeed mimicked the effect of Vangl2/Pk co-injection on Fz, resulting in reduced Fz7 clustering upon Wnt11 induction, formation of cytoplasmic puncta near the plasma membrane, and diminished plasma membrane localization (Suppl. Fig. 13).

To further test whether Vangl2 needs direct interaction with Pk in order to down-regulate Fz7 stability at the plasma membrane and Fz7 patch formation in response to Wnt11, we analyzed Vangl2 R177H variant that specifically reduces Vangl2-Pk interaction (Fig.2 and Suppl. Fig. 6).

We found that unlike wild-type Vangl2, co-injecting Vangl2 R177H with Pk failed to significantly diminish Wnt11-induced Fz7 patch formation or cause cytoplasmic Fz7 puncta (Fig. 5). These data support the notion that direct Pk-Vangl2 interaction is required for efficient sequestration of Dvl from Fz, thereby reducing Fz stability on the plasma membrane.

**Figure 5.**
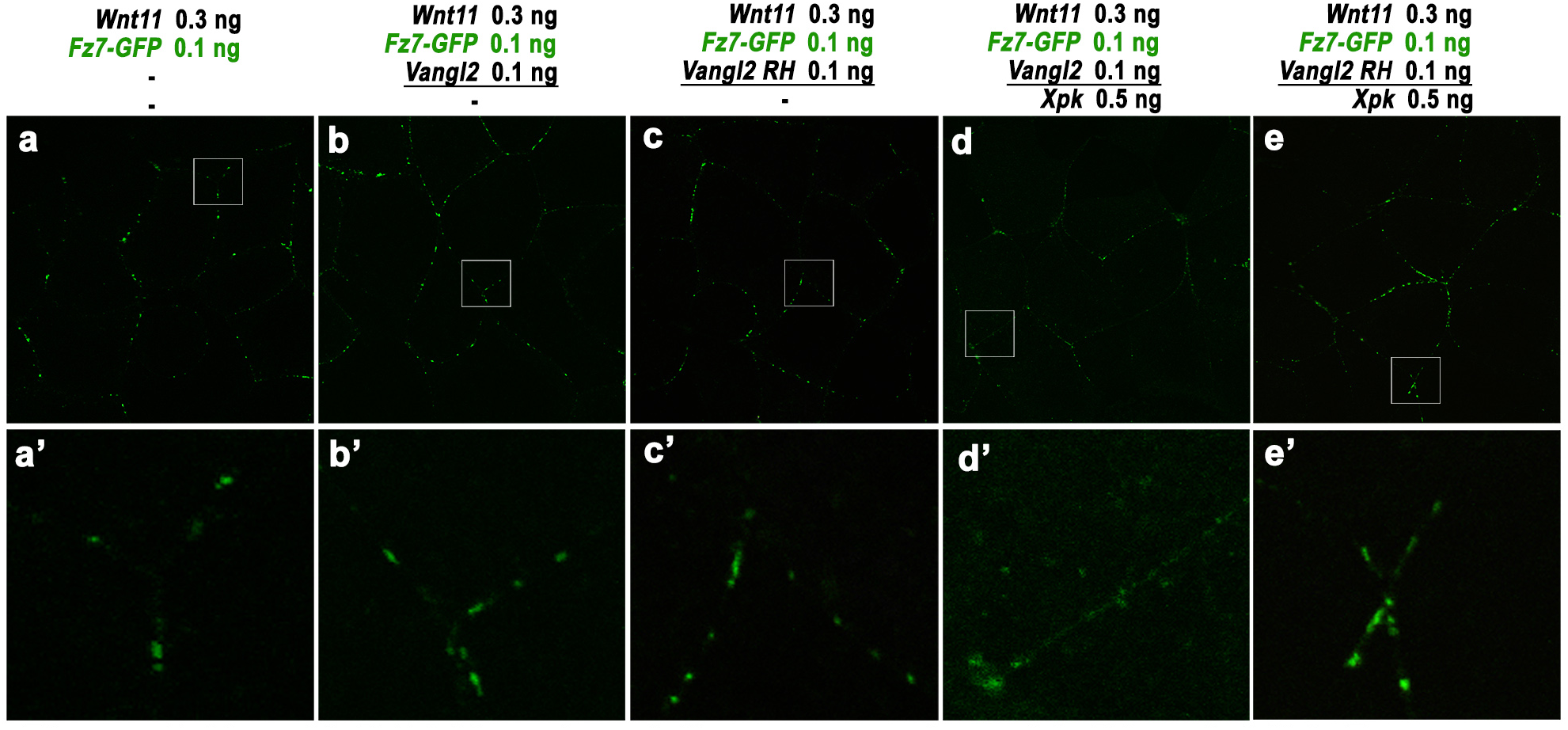
Vangl2 R177H variant fails to synergize with Pk to inhibit Fz patch formation and down-regulate Fz stability at the plasma membrane. Wnt11 induced formation of Fz7-GFP patches on the plasma membrane (a, a’) was not affected by moderate expression either wild-type Vangl2 (b, b’) or Vangl2 R177H variant (c, c’) alone. *Xpk* co-injection synergized with wild-type Vangl2 to diminish Fz7 patch formation and induce cytoplasmic Fz7 puncta (d, d’), but the synergy was not observed with Vangl2 R177H (e, e’).

As a final test for this idea, we examined Dvl phosphorylation known to be inducible by Fz and non-canonical Wnt signaling activation (Axelrod, 2001; Klein et al., 2006; Rothbacher et al., 2000; Shimada et al., 2001; Strutt et al., 2019; Strutt et al., 2006). *Xenopus* extract from embryos injected with *flag-Dvl2* in the DMZ often shows Dvl2 migrating as two bands. The upper, slower migrating band increases in intensity from stage 10 to 12, correlating with the onset and progression of CE (Fig. 6a). The slower migrating form of Dvl2 is increased by Fz7 co-injection, but eliminated by phosphatase treatment (Fig. 6b). Conversely, high level Vangl2 over-expression reduces the phosphorylated form of Dvl2, while Fz7 can counter Vangl2’s effect to increase Dvl2 phosphorylation when co-injected (Fig. 6c). Furthermore, our co-IP experiment demonstrated that only the faster migrating, presumably unphosphorylated form of Dvl2 could be pulled down by Vangl2 in *Xenopus* (Fig. 6d), suggesting that Vangl2-bound Dvl2 may be shielded from Fz-induced phosphorylation. High level Pk (1 ng *Xpk*) is not sufficient to significantly reduce phosphorylated form of Dvl2 when injected alone, but can synergize with moderate level of co-injected Vangl2 to reduce Dvl2 phosphorylation (Fig 6e, f). Together with our previous findings, these data suggest that Pk facilitates Vangl2 to sequester Dvl2 from Fz and, in turn, Fz-induced phosphorylation.

**Figure 6.**
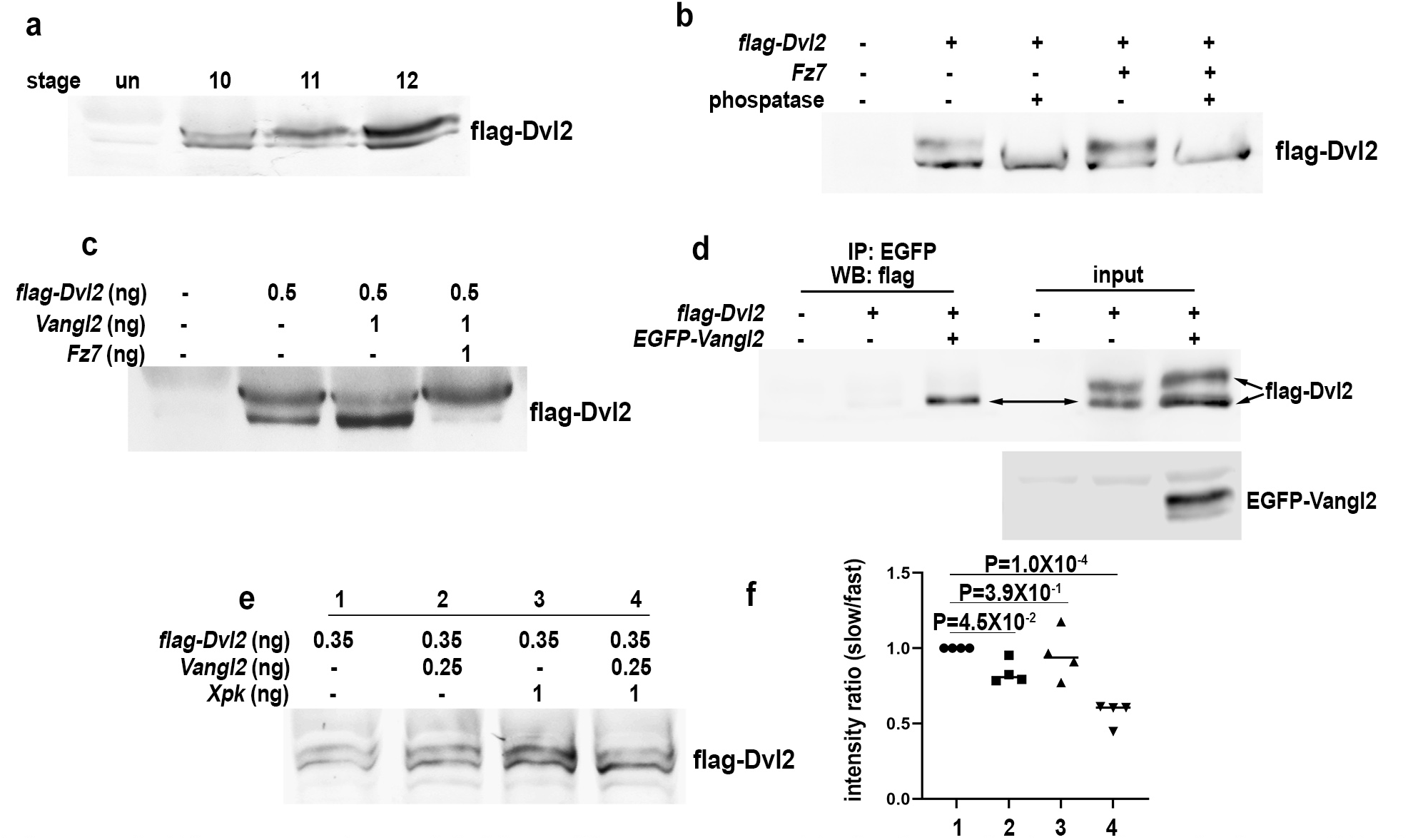
Pk synergize with Vangl2 to prevent Fz-induced phosphorylation of Dvl. In *Xenopus* extract, DMZ injected Dvl2 migrates as two bands, with the slower migrating band increasing in intensity from stage 10 to 12 as CE starts and progresses during gastrulation (a). The slower migrating form of Dvl2 is increased by Fz7 co-injection, but eliminated by phosphatase treatment (b). High level Vangl2 injection can reduce the phosphorylated form of Dvl2, while Fz7 can counter Vangl2’s effect to increase Dvl2 phosphorylation (c). Co-IP experiment indicates that only the faster migrating, presumably unphosphorylated form of Dvl2 can be co-immunoprecipitated by Vangl2 (d). High level *Xpk* (1 ng) or moderate *Vangl2* (0.25 ng) cannot significantly reduce phosphorylated form of Dvl2 when injected individually, but their co-injection can suppress Dvl2 phosphorylation (e). (f) Quantification of the ratio between the slow migrating/ phosphorylated and fast migrating/ unphosphorylated forms of Dvl2 in (e), n=3 biological repeats.

### Ror2 facilitates transition of Dvl2 from Vangl2 to Fz complexes in response to non-canonical Wnt

The above results prompted us to ask that if Dvl is sequestered at the plasma membrane by Vangl2/Pk, how it may transition to form a complex with Fz in response to non-canonical Wnt? As Ror2 has been shown to act as a non-canonical Wnt co-receptor capable of interacting with both Fz and Vangl2 during CE (Gao et al., 2011; Grumolato et al., 2010; Hikasa et al., 2002; Ho et al., 2012; Wallkamm et al., 2014), we hypothesized that Ror2 may be a key component to shuttle Dvl between Vangl2 and Fz.

To test this idea, we first examined the functional relationship between Ror2 and Vangl2. Injecting moderate amount of *Xenopus ror2* mRNA (*Xror2;* 0.05-0.1 ng) can efficiently rescue the severe CE defects induced by 0.2 ng of *Vangl2* mRNA (Suppl. Fig.14a, b), supporting the idea that like Dvl2, Ror2 antagonizes Vangl2 to activate non-canonical Wnt signaling during CE.

Secondly, we tested that at the cellular level, how Wnt11 may induce Ror2 to cluster into patches and how Ror2 patches may correlate with Dvl2 and/or Vangl2 patches. Similar to Fz and Dvl, Ror2-EGFP can be induced to form patches on the plasma membrane by Wnt11 (Fig. 7a-f). Upon co-injection with Dvl2-mCh, the Ror2 patches are overlapped with Dvl2 patches (Fig.7g-i). Close examination of these patches revealed that Ror2, like Fz, accumulates with Dvl2 to high levels in the center of the patches (red arrowhead in Fig. 7g’, j). But unlike Fz, Ror2 patches are slightly longer and extend outside of Dvl2 patches (green arrowheads in Fig. 7g’, j). Quantification indicated that the signal intensity ratio between Ror2 and Dvl2 is increased over two folds at the border of the patches (Fig. 7j, bottom panel). This is reminiscent of Vangl2 enrichment at this region (Fig. 3g’-i’; r), and suggests that Ror2 and Vangl2 may accumulate together at the border of Dvl2 patches to form a complex with diminished amount of Dvl2. Also similar to Vangl2, Ror2 continues to display broad membrane distribution outside of Dvl2 patches (Fig. 7d, g) in the presence of Wnt11, differing from Dvl2 and Fz that are localized exclusively within patches (Fig. 7h,i; Fig.3d-f). These results suggest that at least under the moderate over-expression condition for our imaging experiments, a portion of Ror2 may remain tethered to Vangl2 on the plasma membrane whereas most Dvl2 dissociates from Vangl2 to cluster with Fz in response to Wnt11.

**Figure 7.**
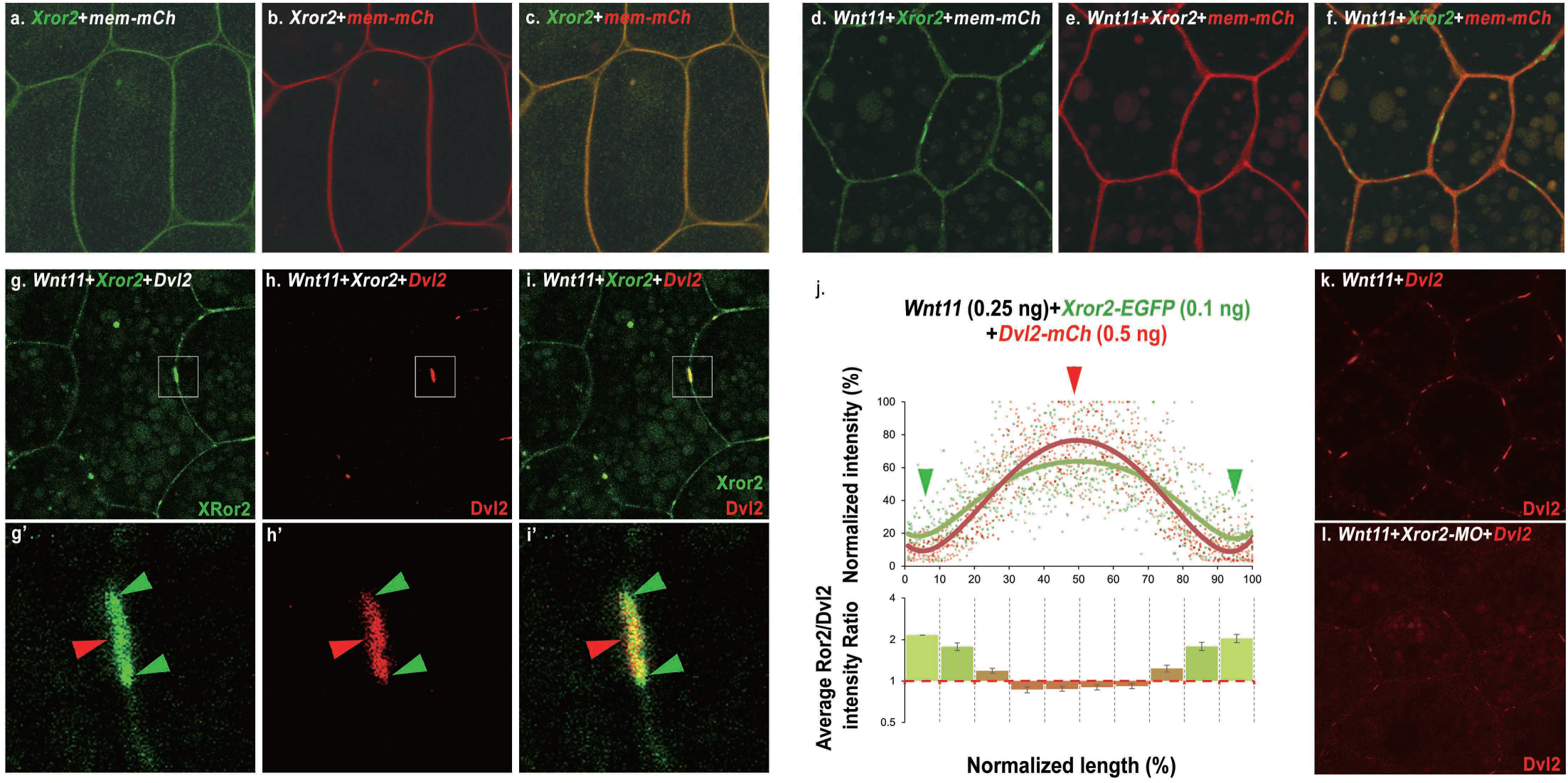
Ror2 is an obligatory component of the Fz/Dvl cluster complex induced by Wnt11. EGFP-tagged *Xenopus* Ror2 (0.1 ng mRNA injection) is distributed homogeneously on the plasma membrane when injected with *membrane-mCherry* (*mem-mCh*, a-c), but can be induced to form distinct patches upon *Wnt11* co-injection (d-f, 0.25 ng). With Dvl2-mCh co-injection (0.5 ng), the Ror2 patches show overlap with Dvl2 patches (g-i). Unlike Dvl2, however, Ror2 additionally displays broad distribution along the entire cell cortex. (compare g to h, and also see (d)). The enlarged views show that both Ror2 and Dvl2 accumulate at the center of the patches (red arrowhead), but Ror2 patches are slightly longer and extend beyond the border of Dvl2 patches (green arrowheads in g’-i’). (j) Measurement of the relative intensity of Dvl2-mCh along the patches with Ror2-EGFP (upper panel), and quantification of ratio between Ror2 and Dvl2 intensity along the patches (bottom panel). Wnt11 induces Dvl2-mCh patch formation on the cell cortex (k) is blocked by co-injecting 25 ng *Xror2* morpholino (*Xror2-*MO) (l).

To test whether Ror2 is required for Dvl2 patch formation in response to Wnt11, we used a verified morpholino to knock down endogenous XRor2 (Schambony and Wedlich, 2007) and found that Dvl2 patch formation is significantly diminished in Xror2 morphants (Fig. 7k, l). Collectively, these data indicate that Ror2, a molecular partner of Vangl2, is an obligatory component of the Fz/Dvl cluster induced by Wnt11.

To further scrutinize the molecular mechanism, we tested biochemically whether Ror2 is required for Dvl2 to dissociate from Vangl2 in response to Wnt11. Unexpectedly, the co-IP data showed that the steady-state binding between Dvl2 and Vangl2 appeared to be reduced by XRor2 knockdown (Supplementary Fig. 15a, a’). The reason for this is unclear, since our imaging showed normal Dvl2 plasma membrane recruitment by Vangl2 with Ror2 knockdown (Supplementary Fig. 16). Irrespective of the reason for the decreased Dvl2-Vangl2 co-IP with XRor2 knockdown, co-injection of Wnt11 cannot further reduce Dvl2-Vangl2 binding in the absence of Ror2 (Supplementary Fig. 15a, a’).

Together, these data support the notion that Ror2 is required for Dvl2 to transition from Vangl2 to Fz in response to Wnt11. To test how Ror2 may facilitate this transition, we performed co-IP and imaging experiments. Our imaging data showed that unlike Vangl2 and Fz7, Ror2 cannot recruit co-injected Dvl2 to the plasma membrane (Suppl. Fig. 17). Also similar to a previous report (Gao et al., 2011), our co-IP experiment detected Ror2 interaction with Vangl2 but not Dvl2 (Suppl. Fig. 15b), suggesting that Ror2 may bind directly to Vangl2 but not Dvl2. As Ror2 and Dvl2 can both bind to Vangl2, we reasoned that they could interact indirectly through their mutual binding with Vangl2. We thus performed fluorescence-detection size exclusion chromatography (FSEC) with protein extract from *Xenopus* embryos injected with *Xror2-EGFP*, *HA-Vangl2* and *Flag-Dvl2*. The elution of Ror2-EGFP was monitored by a fluorescence detector following size exclusion chromatography. The fractions of different molecular size were collected and analyzed by western blot. We found co-fractionation of Ror2, Vangl2 and Dvl2 in fractions 14, 15 and16 (with the approximate molecular weight of 773-1717, 348-773 and 166-348 kD, respectively; Supplementary Fig. 15c). This result supports our hypothesis that Ror2, Vangl2 and Dvl2 form complexes *in vivo*. We envision a model in which an Ror2/Vangl2/Dvl2 complex serves two purposes during CE: it allows Vangl2 to simultaneously sequester both Ror2 and Dvl2 and keeps them inactive, while in response to non-canonical Wnt it enables Ror2 to shuttle Dvl to Fz (See Fig. 9 and Discussion below).

### Bimodal regulation of Ror2 by Vangl2/Pk during non-canonical Wnt signaling

We used Wnt11-induced patch formation as a readout to test the above model. First, we tested whether Vangl2 may synergize with Pk to sequester Ror2 from Fz7 as it does to Dvl2 (Fig. 3 and 4, Suppl. Fig. 9 and 11). When td-Tomato tagged Ror2 and EGFP tagged Fz7 are co-expressed with Wnt11 in either animal cap (Fig. 8) or DMZ (Suppl. Fig. 18) explants, they form clusters that overlap. Moderate over-expression of Vangl2 or XPk individually does not affect Ror2/Fz7 co-clustering within Wnt11-induced patches (compare Fig. 8a-c’ to d-i’, Supplementary Fig. 18a-c’ to d-i’). Vangl2 and Pk co-injection, however, significantly reduced Ror2/Fz7 patches induced by Wnt11 into small puncta (Fig. 8j-l; Supplementary Fig. 18j-l), and caused Fz7 to form intracellular puncta near the plasma membrane (Fig. 8k’, l’, arrows; Supplementary Fig. 178’, l’). Interestingly, like Dvl2 (Fig. 4j’, k’), Ror2 is not present in these Fz7 puncta (compare arrows in Fig. 8j’ to k’ and in Supplementary Fig. 18j’-k’) but remained on the plasma membrane, presumably with Dvl2 and Vangl.

**Figure 8.**
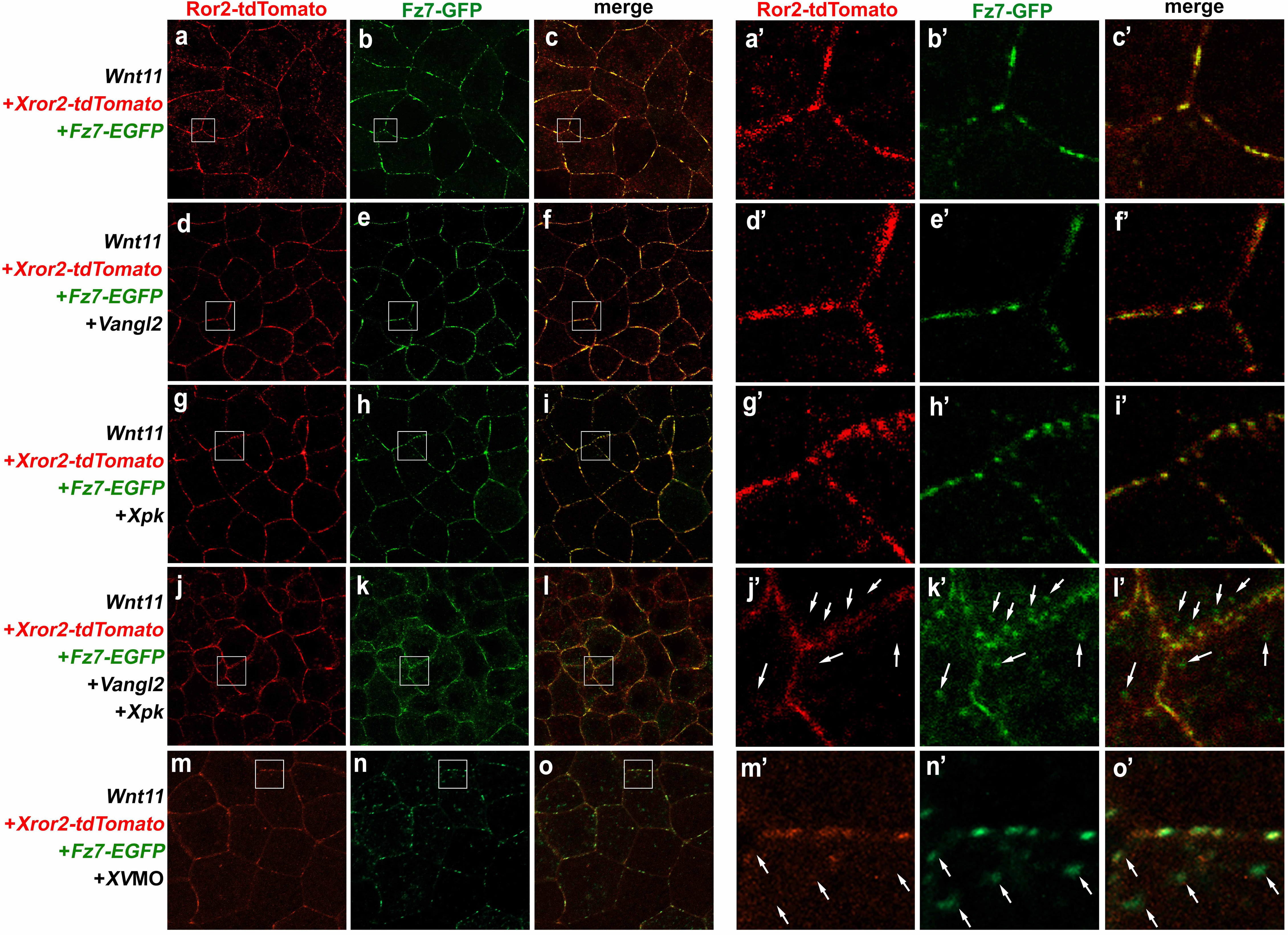
Vangl2/Pk exert bimodal regulation of Ror2 in non-canonical Wnt signaling. (a-c) In animal cap explants, Wnt11 induces co-injected XRor2-tdTomanto and Fz7-EGFP to co-cluster into patches on the cell cortex. Overexpression of moderate level of Vangl2 (d-f, 0.1 ng) or Pk (g-I, 0.5 ng) individually does not perturb co-clustering of Ror2 with Fz7 into patches response to Wnt11, but their co-overexpression (j-l) diminishes Ror2-Fz7 patches into small puncta and cause Fz7 to form cytoplasmic puncta whereas Ror2 remained on the plasma membrane (compare arrows in j’-l’). a’-l’ are enlarged view of a-l, respectively. Conversely, partial knockdown of endogenous *XVangl2* with moderate level of *XV*MO (14 ng) also diminished Ror2/Fz7 patch formation in response to Wnt11 (Fig.8m-o’) with simultaneous formation of intracellular puncta around the plasma membrane which contain Fz7 but not Ror2 (Fig.8m’-o’, arrows).

We then tested whether Vangl2 may also be required for Ror2 to form Wnt11-induced patches with Fz7 since our model predicts that, by bridging Ror2 and Dvl into a complex, Vangl2 helps Ror2 to shuttle Dvl to Fz in response to non-canonical Wnt (Fig. 9). We found that partial knockdown of endogenous *XVangl2* with *XV*MO indeed reduced Ror2/Fz7 patches formed in response to Wnt11 (Fig. 8m-o’) with simultaneous formation of Fz7 intracellular puncta around the plasma membrane (Fig.8m’, n’, arrows), similar to co-overexpression of Vangl2 and XPk (Fig.8k’, l’). Collectively, these data support our model and suggest that with Pk, Vangl2 exerts bimodal regulation of Ror2 in non-canonical Wnt signaling.

**Figure 9.**
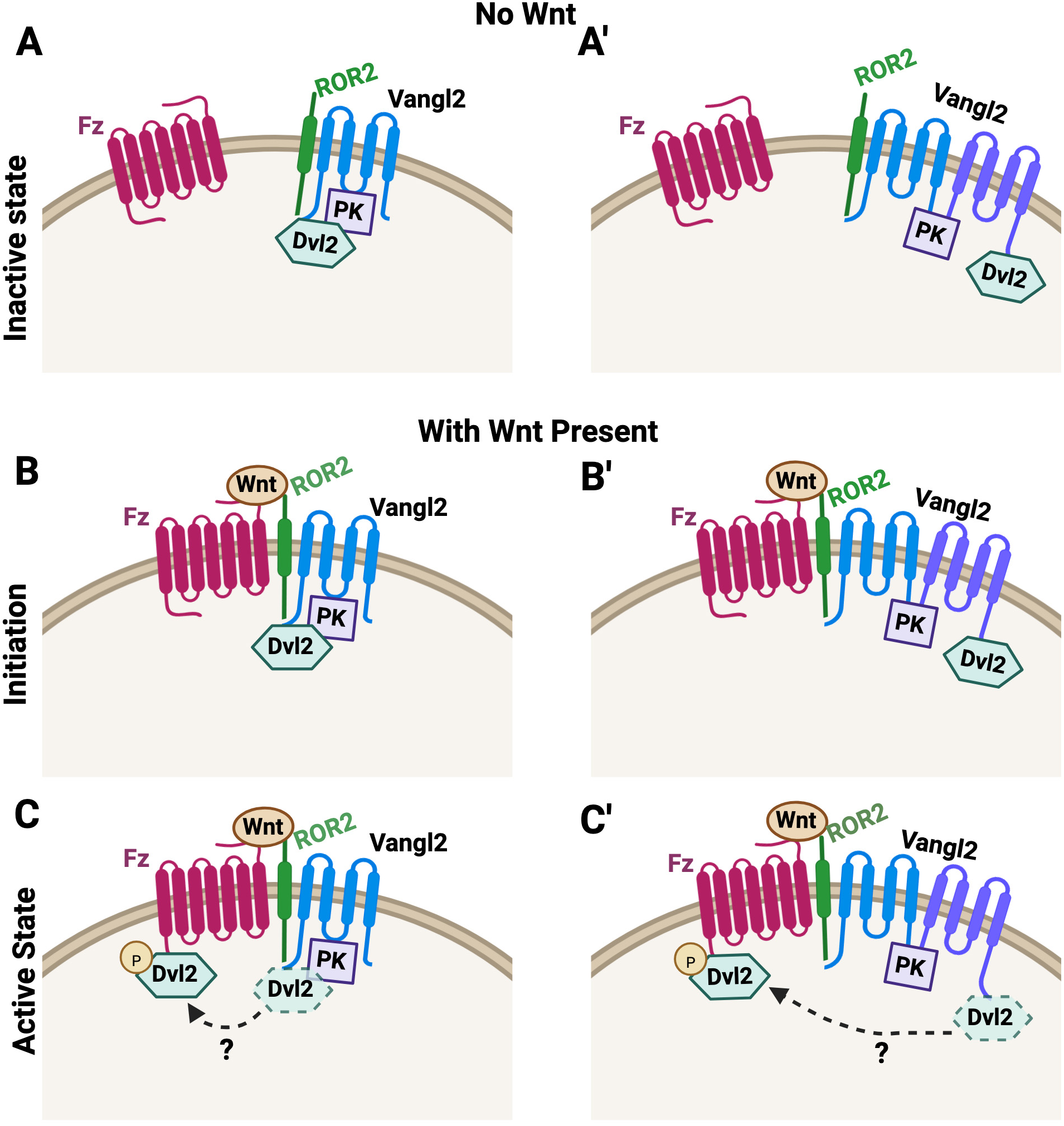
An integrated model for non-canonical Wnt signaling regulation during CE. (a, a’) In the absence of non-canonical Wnt, Pk helps Vangl to act as an adaptor that brings together Dvl and Ror, through either simultaneous binding of both Dvl and Ror to a single Vangl (a) or oligomerization of Vangl proteins bound separately to Dvl and Ror (a’), and keeps both Dvl and Ror inactive to prevent ectopic non-canonical Wnt signaling. (b, b’) Non-canonical Wnt initiates signaling by triggering Fz-Ror heterodimerization, and in turn, the complexes consisting of Ror/Vangl/Dvl are brought close to Fz to deliver Dvl for activation of downstream targets. (c, c’) Non-canonical Wnt also induces other events such as Dvl phosphorylation to facilitate Dvl dissociation from Vangl and transition to Fz in a spatially and temporally controlled manner.

## Discussion

Early fly studies identified six proteins that act as core members to coordinate cellular polarity across the plane of the epithelium. In-depth genetic, biochemical and imaging studies have subsequently elucidated how the six core PCP proteins interact within and between cells to establish feedback loops that partition Vang/Pk and Fz/Dsh/Dgo clusters on opposing cell cortexes to coordinate polarity (Amonlirdviman et al., 2005; Goodrich and Strutt, 2011). These studies establish a foundation to understand the action of PCP proteins in static epithelial cells. They do not, however, seem to provide direct explanation for how PCP proteins regulate polarized and dynamic cell behavior during CE, where asymmetric partitioning of core PCP proteins has not been consistently observed and non-core PCP proteins, including non-canonical Wnt ligands, co-receptors Ror1/2 and a cytoplasmic protein Dact1, are also critically involved. Furthermore, adopting core PCP proteins to regulate CE is likely a vertebrate-specific adaption during evolution, since fly germband extension, a CE-like morphogenetic event, does not involve core PCP proteins (Zallen and Wieschaus, 2004). We previously provided evidence for a model that during CE, Vangl exerts bimodal regulation of Dvl by cell-autonomously recruiting Dvl to the plasma membrane in an inactive state, and simultaneously poising Dvl for activation upon binding of Fz to non-canonical Wnt ligands (Seo et al., 2017). Our recent work tested this model by studying how Dact1, a vertebrate specific protein, modulates Dvl-Vangl interaction during non-canonical Wnt signaling and CE (Angermeier et al., 2025). In the current study, we further tested this model and demonstrated that Pk functionally synergizes with Vangl2 to inhibit Dvl2 during CE in *Xenopus*. Mechanistically, Pk binding to Vangl2 helps Vangl2 to sequester Dvl and constrain its transition to Fz. Moreover, Pk seems to play a similar role in assisting Vangl2 to sequester Ror2, whereas Ror2 is required for Dvl2 to transition from Vangl to Fz in response to non-canonical Wnt. We propose an updated model for the bimodal regulation in which Vangl2/Pk bring both Dvl2 and Ror2 into an inactive complex that prevents ectopic non-canonical Wnt signaling. On the other hand, this complex can also be coupled with Fz upon non-canonical Wnt induced binding between Ror2 and Fz, delivering Dvl to Fz to initiate non-canonical Wnt signaling (Fig. 9). Therefore, Vangl mediated plasma membrane recruitment of Dvl and pre-assembly of Dvl/ Vangl/ Ror complex can also accelerate non-canonical Wnt signaling activation by facilitating Dvl presentation to ligand-bound Fz. Our model provides a new framework to decipher how core PCP proteins are integrated with non-core proteins to tightly control the threshold and dynamics of non-canonical Wnt/ PCP signaling during CE.

### Regulation of Vangl-Dvl interaction by Pk to suppress non-canonical Wnt signaling

Our previous work proposed that Vangl-Dvl interaction provides a key switch to the central logic of non-canonical Wnt siganling by enriching Dvl around the plasma membrane for effective access to Fz, while at the same time keeping Dvl inactive to prevent ectopic signaling (Seo et al., 2017). In the current study, we found that the Vangl2 R177H variant (Kibar et al., 2011), which can traffick properly and bind to and recruit Dvl2 to the plasma membrane like wild-type Vangl2 (Fig. 2, Suppl. Fig. 5), is less capable of inhibiting CE and rescuing Fz/Dvl over-expression induced CE defect (Suppl. Fig.5). The results suggest that binding to Dvl *per se* is not sufficient for Vangl2 to suppress Dvl during CE. Interestingly, Vangl2 R177H displays significantly reduced binding and functional synergy with Pk. We therefore reason that interaction with Pk is necessary for Vangl2 to efficiently sequester Dvl from Fz or downstream targets like Daam1. Our biochemical and imaging experiments provide supporting evidence to this idea (Fig. 3-5; Suppl. Fig. 8, 9, 11, 16).

There are at least three possibilities accounting for how Pk can assist Vangl2 to sequester Dvl. First, given that Dvl, Pk and Vang/Vangl can mutually interact with each other (Bastock et al., 2003; Humphries et al., 2023; Jenny et al., 2003; Takeuchi et al., 2003; Tree et al., 2002), Pk may stabilize a ternary Dvl/Pk/Vangl complex by simultaneously interacting with both Vangl and Dvl (Fig. 9a). We, however, do not favor this possibility because 1) the binding between Pk and Dvl was reported to be quite weak (Bastock et al., 2003); 2) our unpublished data show that ΔPL, a Pk mutant lacking the PET/LIM domains necessary for Dvl binding (Takeuchi et al., 2003), can largely mimic wild-type Pk function. Furthermore, a recent study in flies showed that the phosphorylation status of a conserved tyrosine in the cytoplasmic tail of Vang provides opposite binding preference for Pk and Dsh, suggesting that simultaneous binding of both Pk and Dsh to the C-terminus of Vang may not be possible (Humphries et al., 2023).

Secondly, Pk may regulate biochemical modification on Vangl2 to strengthen Vangl2-Dvl interaction. The best known modification on Vang/Vangl is phosphorylation at several N-terminal serine/ threonine residues in response to Wnt/ Fz (Gao et al., 2011; Kelly et al., 2016; Yang et al., 2017). Whereas one study in flies reported that Pk can prevent Vang phosphorylation at these residues to decrease Vang turnover (Strutt et al., 2019), another study in fly S2 cells found that Pk over-expression does not alter Vang N-terminal phosphorylation (Kelly et al., 2016). In our current studies in *Xenopus,* we have not been able to detect Vangl2 phosphorylation consistently, but this possibility remains an interesting idea and should be tested in the future using the reported phosphomutant and phosphomimetic Vangl2 (Yang et al., 2017). Finally, phosphorylation of a conserved C-terminal tyrosine residue of Vang was recently reported to decrease its binding with Dsh (Humphries et al., 2023), and could therefore provide a mechanism to control Vangl-Dvl interaction. But the regulatory mechanism of this tyrosine phosphorylation is not clear and does not seem to depend on Fz. Further studies will be needed to elucidate the role of this tyrosine phosphorylation in vertebrate CE.

Thirdly, Pk binding may induce allosteric change or clustering of Vangl to increase the overall avidity for Dvl binding. Vang/Vangl was proposed to dimerize, and possibly oligomerize into larger cluster through their C-terminal tail and/or transmembrane domains (Belotti et al., 2012; Jenny et al., 2003), and two recent cryo-EM studies revealed that Vangl1/2 can oligomerize into trimers (Song et al., 2025; Zhang et al., 2025). Interestingly, quantitative imaging studies in flies have revealed that in stable PCP clusters, the ratio between Vang and Pk is 6:1 (Strutt et al., 2016). In light of our data suggesting that the intracellular loop between TM2 and 3 in Vangl2 may impact Pk binding in addition to the canonical Pk binding domain at the C-terminal tail (Fig. 2; (Bastock et al., 2003; Humphries et al., 2023; Jenny et al., 2003), it is tempting to speculate that Pk may nucleate or stabilize Vangl oligomer formation through multimeric interactions with different domains on multiple Vangl proteins. Such oligomeric Vangl cluster may form a “cage” to more effectively sequester Dvl due to increased local concentration and/or higher binding affinity resulted from conformational change upon oligomerization or Pk binding.

Our above model seemingly contradicts with the fly studies showing that Vang/Pk clusters are partitioned to the opposite cell cortexes from Fz/Dsh clusters and are clearly devoid of Dsh. These segregated clusters, however, seem to form progressively from initial symmetrically distributed PCP proteins along cell-cell junctions at early stages where Vang does co-mingle with Dsh (Bastock et al., 2003), and a new study further implicated the functional importance of Vang-Dsh binding in fly PCP establishment (Humphries et al., 2023). Persistent contact and stable junctions between neighboring cells may facilitate feedback interaction to partition Vang/Pk from Dsh/Fz (Stahley et al., 2021). In dynamically moving cells during CE, intercellular feedback interactions are likely limited and transient, therefore posing challenges for stable segregation of distinct PCP clusters. Conversely, non-canonical Wnt ligands play a key role during vertebrate CE but not in fly PCP establishment. These differences may lead to some changes in the molecular actions of core PCP proteins (see below).

### An Ror-dependent relay mechanism to deliver Dvl for non-canonical Wnt signaling

The premise of our model is that during CE, Vangl acts via a relay mechanism to first bring Dvl to the plasma membrane, and then releases Dvl to Fz. Pk may tighten up this relay mechanism, via regulating Vangl-Dvl interaction, to increase the efficiency of Dvl plasma membrane recruitment and the threshold at which Dvl can be released to Fz. While the detailed mechanisms for how Dvl can be released from Vangl and transitioned to Fz are yet to be elucidated in further details in the future, our studies identified several factors that contribute to the transition: non-canonical Wnt, the co-receptor Ror2 and Dvl phosphorylation.

Our co-IP and imaging studies showed that Wnt11 can trigger dissociation of Dvl2 from Vangl2 (Fig. 3g-i’; Suppl. Fig. 8 and 9g-i’; (Angermeier et al., 2025; Seo et al., 2017)) and formation of Fz7-Dvl2 clusters at cell-cell contact (Fig. 3d-f’, Suppl. Fig. 9d-f’), indicating that non-canonical Wnt can act extracellularly to trigger the transition of Dvl from Vangl to Fz. It is possible that Wnt binding to Fz can directly induce events in favor of Fz-Dvl association. We, however, also consider the alternate possibility that Wnt binding to the co-receptors Ror1/2 to bring Dvl to Fz.

Like Fz, Ror1/2 also harbor the extracellular cysteine-rich domains known to interact with Wnts and have been shown to heterodimerize with Fz in response to non-canonical Wnt binding (Griffiths et al., 2024; Grumolato et al., 2010). At the same time, like Dvl, Ror2 was reported to bind directly with Vangl2 (Gao et al., 2011). We therefore postulate that Ror2 may shuttle between Vangl and Fz to deliver Dvl in a Wnt dependent manner. We note several intriguing links between Ror2 and Dvl2: 1) they both bind to Vangl2 yet display functional antagonism against Vangl2 during CE in over-expression assays (Fig. 1; Suppl. Fig. 14); 2) they both cluster with Fz in response to Wnt11 (Fig. 3, 7, 8; Suppl. Fig. 9, 16), and importantly Ror2 is required for Dvl2 to dissociate from Vangl2 and cluster with Fz in response to Wnt11 (Fig. 7l; Suppl. Fig. 15a, a’). While Ror2 does not seem to bind Dvl directly in our experiment, they both interact with Vangl2 and our SEC data show that Ror2, Vangl2 and Dvl2 co-fractionate (Suppl. Fig. 15), suggesting that they may form complexes together. Therefore, we envision a model where Vangl acting as an adaptor to bring together Dvl and Ror, either through simultaneous binding of both Dvl and Ror to a single Vangl (Fig. 9a) or self-oligomerization of Vangl proteins bound separately to Dvl and Ror (Fig. 9b). When non-canonical Wnt induces Fz-Ror to heterodimerize, the complexes consisting of Ror/Vangl/Dvl can be brought close to Fz to deliver Dvl and initiate non-canonical Wnt signaling (Fig. 9a’,b’).

In this model, Vangl acts as an unconventional adaptor to simultaneously serve two critical functions: it pre-assembles Ror and Dvl into complexes at the plasma membrane ready to initiate non-canonical Wnt signaling, but at the same time keeps both inactive to prevent ectopic signaling activation. We previously demonstrated that Vangl2 can prevent Dvl from interacting with its downstream effector Daam1 (Seo et al., 2017), and our data in this study suggest that Vangl/Pk may act together to sequesters both Dvl and Ror from Fz as well (Fig. 4 and 8; Suppl. Fig. 11, 16). Based on this model, Vangl2 over-expression will exert excessive suppression to prevent non-canonical Wnt signaling during CE, which can be overcome by co-overexpressing Dvl2 or Ror2 (Fig. 1; Suppl. Fig. 14). Conversely, reducing the dosage of endogenous Vangl2 may decrease the assembly of Ror/Vangl/Dvl complexes, compromising signaling activation in response to non-canonical Wnt (Fig. 8m-o’). Therefore, our model can explain how partial loss of Vangl2 can synergize with loss of positive non-canonical Wnt signaling regulators, including Ror2, Dvl2 and Wnt5a, to cause various severe CE defects reported in the literature (Gao et al., 2011; Qian et al., 2007; Sinha et al., 2012; Wang et al., 2011; Wang et al., 2006). We acknowledge, however, that our model explains primarily the potential molecular actions underlying the regulation of CE at the tissue and genetic levels. Whether and how our model may explain the cellular behavior during CE, such as polarized remodeling of cell junction or extension of cell protrusions, will require further study.

Lastly, our data implicated an intriguing role for Dvl phosphorylation in the transition from Vangl to Fz. We found that flag-Dvl2 phosphorylation is increased as CE progresses in *Xenopus* and can be elevated by Fz7 but suppressed by Vangl2 (Fig. 6b, c). In agreement with our over-expression data in *Xenopus*, loss of both Vangl1 and 2 leads to increased Dvl2 and 3 phosphorylation in cell culture (Mentink et al., 2018). Intriguingly, Vangl2 seems to bind only to unphosphorylated form Dvl2 (Fig. 6d). These observations suggest that either Dvl phosphorylation *per se* or another associated modification can be used as a mechanism to decouple Dvl from Vangl. In support of this view, the basic region and PDZ domain of Dsh/Dvl, which mediates Dvl-Vangl interaction (Angermeier et al., 2025; Park and Moon, 2002), is a strong target of CK1 during PCP signaling in flies (Klein et al., 2006; Strutt et al., 2019; Strutt et al., 2006). Our recent work further revealed that Dvl oligomerization promotes its dissociation from Vangl possibly by occluding the PDZ domain, and Dvl mutants defective at oligomerization also fail to undergo phosphorylation (Angermeier et al., 2025).

Taken together, we propose the second piece of our model (Fig. 9c, c’) that during CE, non-canonical Wnt triggers association of Ror1/2 and Fz to simultaneously accomplish two events: 1) bringing Vangl-sequestered Dvl close to Fz; and 2) activating CK1 or other kinases to phosphorylate Dvl. The combined effects lead to Dvl dissociation from Vangl and transition to Fz in a spatially and temporally controlled manner. On the other hand, by assisting Vangl to sequester Dvl, Pk may suppress the noise from basal CK1 activity, and allow cells to respond more specifically and dynamically to non-canonical Wnt singling during CE.

## Materials and Methods

Animal experiments were performed in agreement with the National Institutes of Health. *Xenopus laevis* adults were maintained according to the established protocols by the Institutional Animal Care and Use Committee at the University of Alabama at Birmingham, under Animal Project Number IACUC-22388. There is no evidence for sexual dimorphism in gastrulating *Xenopus* embryos, which were used as the model organism in the study, so sex was not considered as a biological variable in the study design.

### *Xenopus* embryo manipulation and animal cap/DMZ explants

Embryos were acquired by superovulation, maintained in 0.1% MMR solution until the stages for microinjection. Morpholinos or *in vitro* synthesized RNAs were injected into either the animal side of two-cell-stage embryos or the DMZ of four-cell-stage embryos. For phenotypic analysis, the DMZ-injected embryos were fixed at tailbud stages, and the dorsal view of embryos was captured using a Leica DFC 490 camera mounted on a Leica M205 FCA stereomicroscope. A Fiji macro was utilized to process the images and obtain the projected area and the length of each embryo. Briefly, the macro first extracts individual embryos from the images, measures the area of each embryo, and generates the smallest rectangle that fully encloses each embryo. The length of the rectangle is considered the length of the embryo, while the width is calculated by dividing the embryo’s area by its length. The LWR is then calculated in Excel. The length of the embryos with significant curved shape was correct manually by drawing a line along the anteroposterior axis to measure the maximal distance from the head to the tail of each embryo using the Leica LAS software with Interactive Measurement module. For animal cap elongation assay, ectodermal explants were isolated at stages 9-10 and incubated in 0.5 MMR solution containing 10 ng/ml of Activin B (R&D cat# 659-AB-005). The CE phenotype was quantified by measuring the length of the resulting explants. For fluorescent imaging to determine protein localization, DMZ or animal cap explants from injected embryos were isolated at stage 10-10.5, coversliped and subjected to confocal imaging analysis as described (Angermeier et al., 2025; Seo et al., 2017).

### Co-immunoprecipitation and western blot

RNAs or morpholinos injected embryo or explants were lysed as described for biochemistry experiments (Angermeier et al., 2025; Seo et al., 2017; Tien et al., 2015). For co-immunoprecipitation assay, protein lysates were subjected to pull-down with anti-flag (Sigma Anti-FLAG M2 Magnetic Beads (Cat# 8823)) or anti-Myc antibodies (Pierce Cat.#88843), in a buffer containing 50mM Tris (pH 7.5), 150mM NaCl, 1mM EDTA, 10% Glycerol, 0.5% Triton-x100 and 1x protease inhibitor (Promega #G6521). Western blot detection of proteins was carried out with anti-GFP antibody (Santa Cruz Biotechnology GFP (B-2) (Cat# sc-9996)), anti-myc antibody (Santa Cruz Biotechnology myc Antibody(G-4) (Cat# sc-373712)), anti-Dvl2 (CST Cat.# 3224), anti-XRor2 (Developmental Studies Hybridoma Bank) or anti-flag antibody (Sigma Anti-flag M2 antibody (Cat# F1804)). We followed the protocol by Burckhardt et. al. (Burckhardt et al., 2021) and used FIJI to quantify the relative amount of co-IP. Briefly, the intensity values from the co-IP of any protein was divided by that from the IP of its presumptive partner under each control or experimental condition; the mean ratios from the controls was then used to normalize the ratios under each experimental condition to determine the “Relative Co-IP amount”.

### Fluorescence-detection size-exclusion chromatography (FSEC)

2-cell stage *Xenopus* embryos were co-injected with 3 ng Ror2-EGFP, 0.5 ng Flag-Dvl2 and 1 ng HA-Vangl2 mRNA to the animal side. Animal caps were dissected around stage 10 and cultured in 0.5 X MMR at 15 ℃ overnight. Next day 35 – 40 animal caps were lysed on ice with the 200 mL lysis buffer ( 50 mM Tris pH 7.5, 150 mM NaCl, 0.3% Dodecylmaltoside and Protease inhibitor (Pierce^TM^ Protease Inhibitor)). After centrifuged at 4 ℃ 14000g for 15 min, the supernatant was collected and filtered with 0.22 mm syringe filter. 100 mL lysate was loaded onto Superose 6 Increase 10/300 GL column (Cytiva, Marlborough, MA) pre-equilibrated in SEC buffer (Tris 50 mM, pH 7.5, NaCl 150 mM, Dodecylmaltoside 0.03%). The eluted Ror2-GFP was monitored by RF-10AX fluorescence detector (Shimadzu, Japan) following size-exclusion chromatography. Fractions were collected and concentrated (Pierce^TM^ concentrator PES 10K MWCO), and used for Western blot.

### Imaging and analyses

For imaging, 0.1-0.5 ng of mRNA encoding Dvl2-mCh, Dvl2-mSc, Dvl2-EGFP, Fz7-EGFP, XRor2-tdTomato, XRor2-EGFP, EGFP-Vangl2, EGFP-Vangl2 RH, flag-XPk, EGFP-mPk2, XWnt11, mem-GFP and mem-mCh were injected in various combination into the animal regions at the two-cell stage and dissected at ∼St.9, or the DMZ at the four-cell stage and dissected at St. 10.25. Alternatively, *flag-Xpk* injected animal caps were dissected and fixed in 4% PFA for immunofluorescence staining with an anti-flag antibody. Dissected animal cap or DMZ explants were imaged on an Olympus FV1000 with a 20x water immersion objective or a Zeiss LSM 900 equipped with Airyscan2 and a 20x air objective. Images were imported into ImageJ (NIH). Images from at least three different embryos collected on different days were analyzed per injection group.

The relative protein level between the plasma membrane and cytoplasm was quantified by comparing the fluorescent intensity. Briefly, regions corresponding to the plasma membrane and cytoplasm were defined, and the average fluorescence intensity within each region was measured using ImageJ. The ratio of plasma membrane to cytoplasmic fluorescence intensity was then calculated.

Protein enrichment pattern within clusters was analyzed using ImageJ. Briefly, the fluorescence intensity of each pixel along a defined cluster was measured. To account for differences in cluster size, pixels were sequentially numbered from one end of the cluster to the other. The total pixel count for each cluster was normalized to 100, and each pixel was assigned a relative position expressed as a percentile. Similarly, fluorescence intensity was normalized by setting the highest pixel intensity within each cluster to 100, and expressing all other pixel intensities as percentiles of that maximum. A scatter plot was generated with the x-axis representing the relative position and the y-axis representing the normalized fluorescence intensity of each pixel. The regression curve was then fitted to the data to characterize the distribution pattern of fluorescence intensity across the cluster. 10 clusters from 5-10 embryos were used for statistical analysis.

To analyze Fz7 endocytosis, vitelline membrane of injected embryos was removed at stage 10.5, and embryos were incubated in 0.1X MMR containing 5ug/ml FM4-64FX (Thermo Fisher, Cat# F34653) for 30min at room temperature before dissection in 0.1X MMR. Dissected DMZ explants were coversliped, and images were captured without fixation. Images from at least three different embryos collected on different days were analyzed per injection group.

### RNAs and morpholinos

XWnt11, EGFP-Vangl2, HA-Vangl2, GFP-XFz7, flag-XFz7, tdT-tomatoRor2, myc-Vangl2, Dvl2-mCherry, Dvl2-flag, GFP-mPk2, flag-XPk, flag-PL, flag-ΔPL, were transcribed in vitro using mMESSAGE mMACHINE SP6 Transcription Kit (Ambion cat#1340). Xenopus Vangl2-morpholino (XVMO), Ror2-morpholino (Xror2-MO) and Pk-morpholino (XPkMO) are the same as previously described (Darken et al., 2002; Schambony and Wedlich, 2007; Takeuchi et al., 2003). The dosage of each RNA or morpholino is described in each figure.

## ACKNOWLEDGEMENTS

We thank Dr. Naoto Ueno for providing the flag-XPk construct. The anti-XRor2 monoclonal antibody was obtained from the Developmental Studies Hybridoma Bank, created by the NICHD of the NIH and maintained at The University of Iowa, Department of Biology, Iowa City, IA 52242.

## COMPETING INTERESTS

The authors declare no competing or financial interests.

## FUNDING

This work was supported by grants R35GM131914 (J.D.A.), GM127371 (C.C.), and HL109130, HL138470 and AR081646 (J.W.) from the National Institutes of Health.

## Supplementary Figures

**Supplementary Figure 1.**
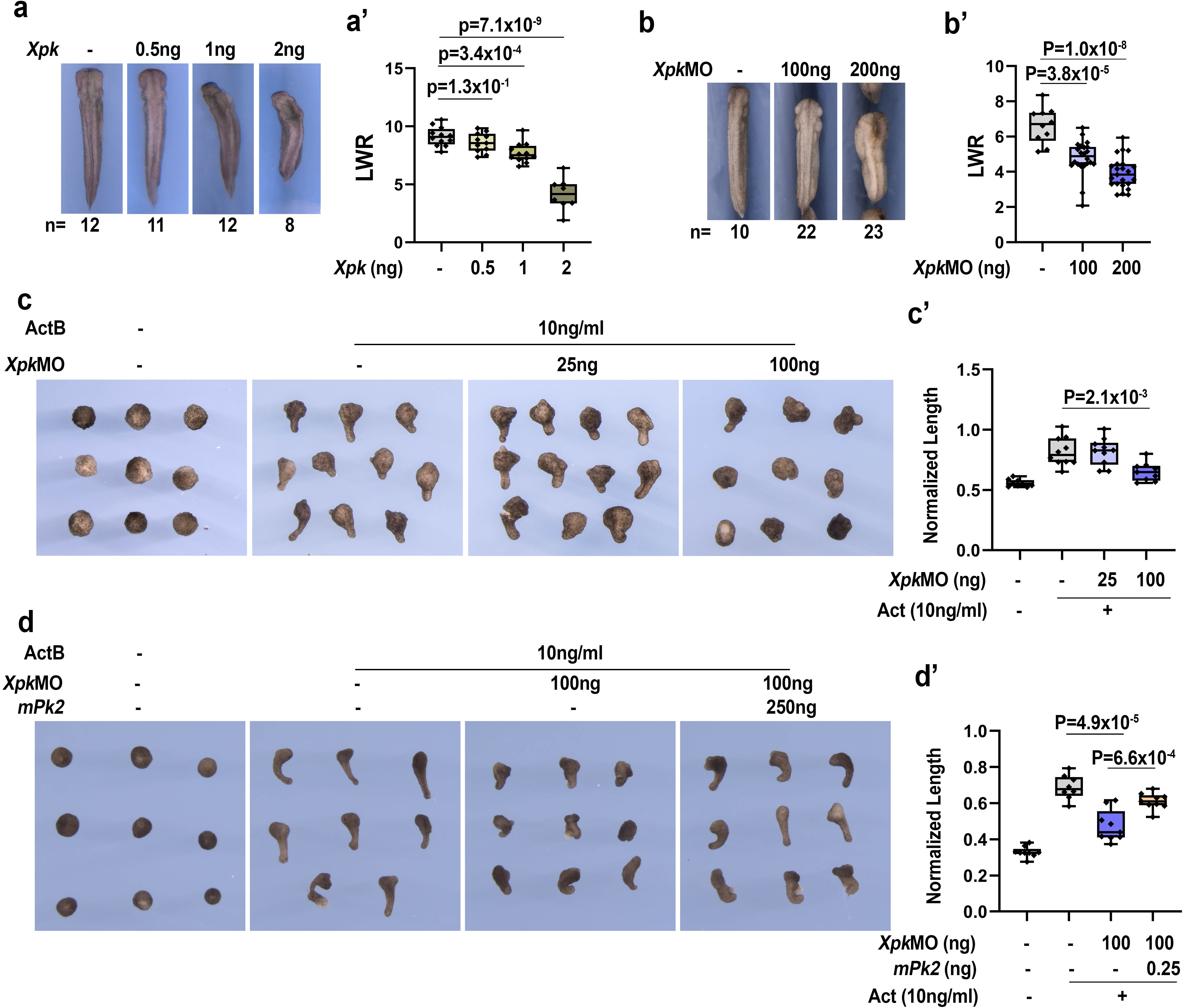
DMZ injection to over-express (a, a’) or morpholino knockdown (b, b’) of *Xpk* can dose-dependently block CE, resulting in shorten body axes and reduced LWR. In the animal cap explant, *Xpk*MO knockdown can also block activin induced CE (c, c’), but the defect can be rescued by co-injecting RNA encoding mouse Pk2 (d, d’). CE phenotype in (a) and (b) was determined by quantifying the length-to-width ratio (LWR) of the embryos in each group. Experiments were repeated three times and the total number of embryos analyzed is indicated below each panel in (a) and (b). Quantification of the length of the animal cap explants are in (c’) and (d’). Data are presented as box plots in (a’), (b’), (c’) and (d’), with the whiskers indicating the minima and maxima, the center lines representing median, the box upper and lower bounds representing 75th and 25th percentile, respectively. Two-tailed, unpaired T-test was used to compare the LWR of different groups, and the p vales are indicated in (a’)-(d’) between different groups.

**Supplementary Figure 2.**
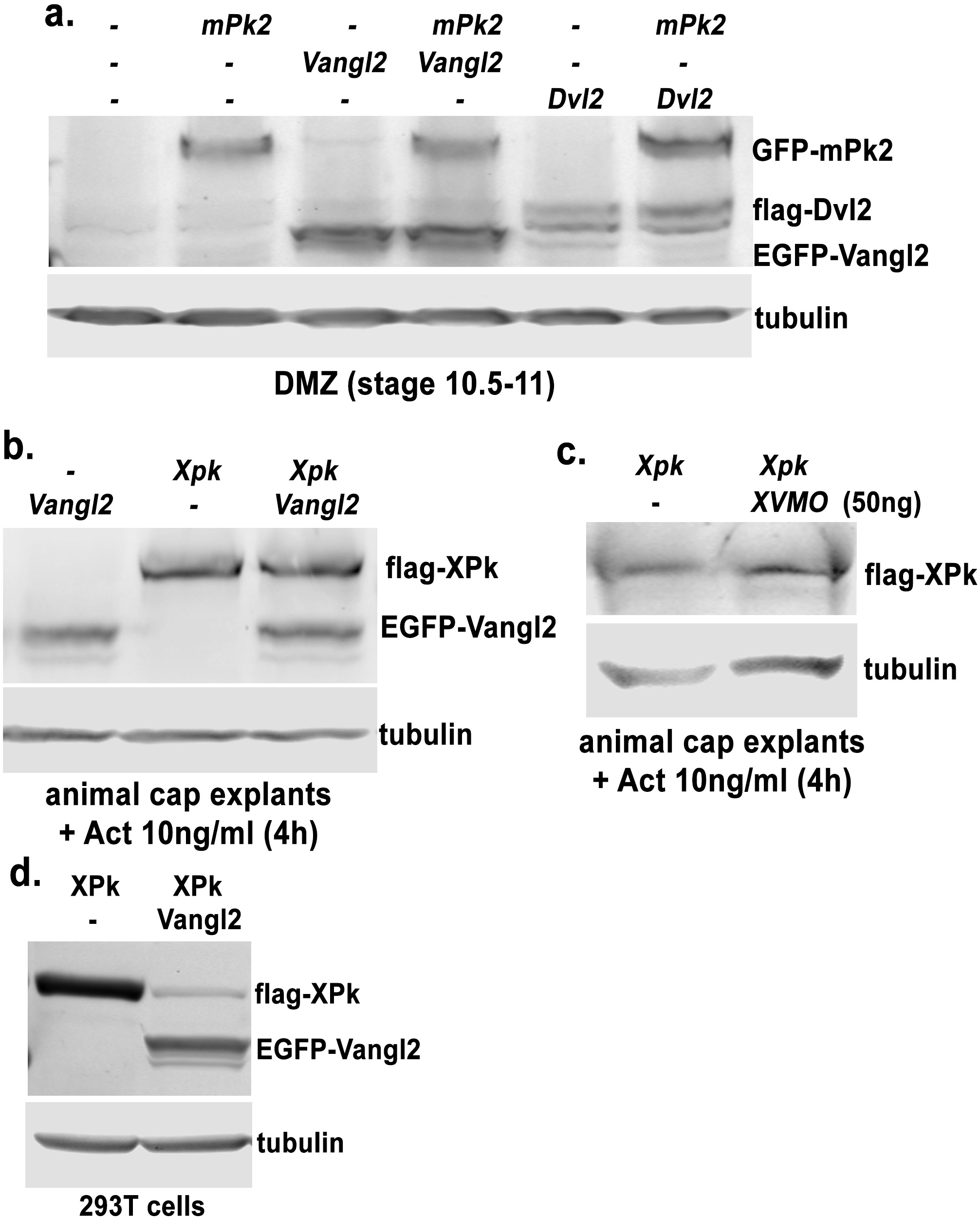
*EGFP-Vangl2*, *flag-Dvl*2 and *GFP-mPk2* mRNA were injected into the DMZ, either alone or together, at 4 cell stage, and embryos were harvested at stages 10.5-11 for western blot. Co-overexpressing Vangl2 with mPk2, or Dvl2 with mPk2, does not significantly alter each other’s protein level (a). Similarly, co-overexpressing Flag-XPk with EGFP-Vangl2 in activin treated animal cap explant also does not alter each other’s protein level (b). Knocking down endogenous *Xvangl2* does not alter the protein level of injected Flag-XPk either (c). Interestingly, however, co-transfecting Vangl2 with XPk into 293T cells reduced XPk protein level (d).

**Supplementary Figure 3.**
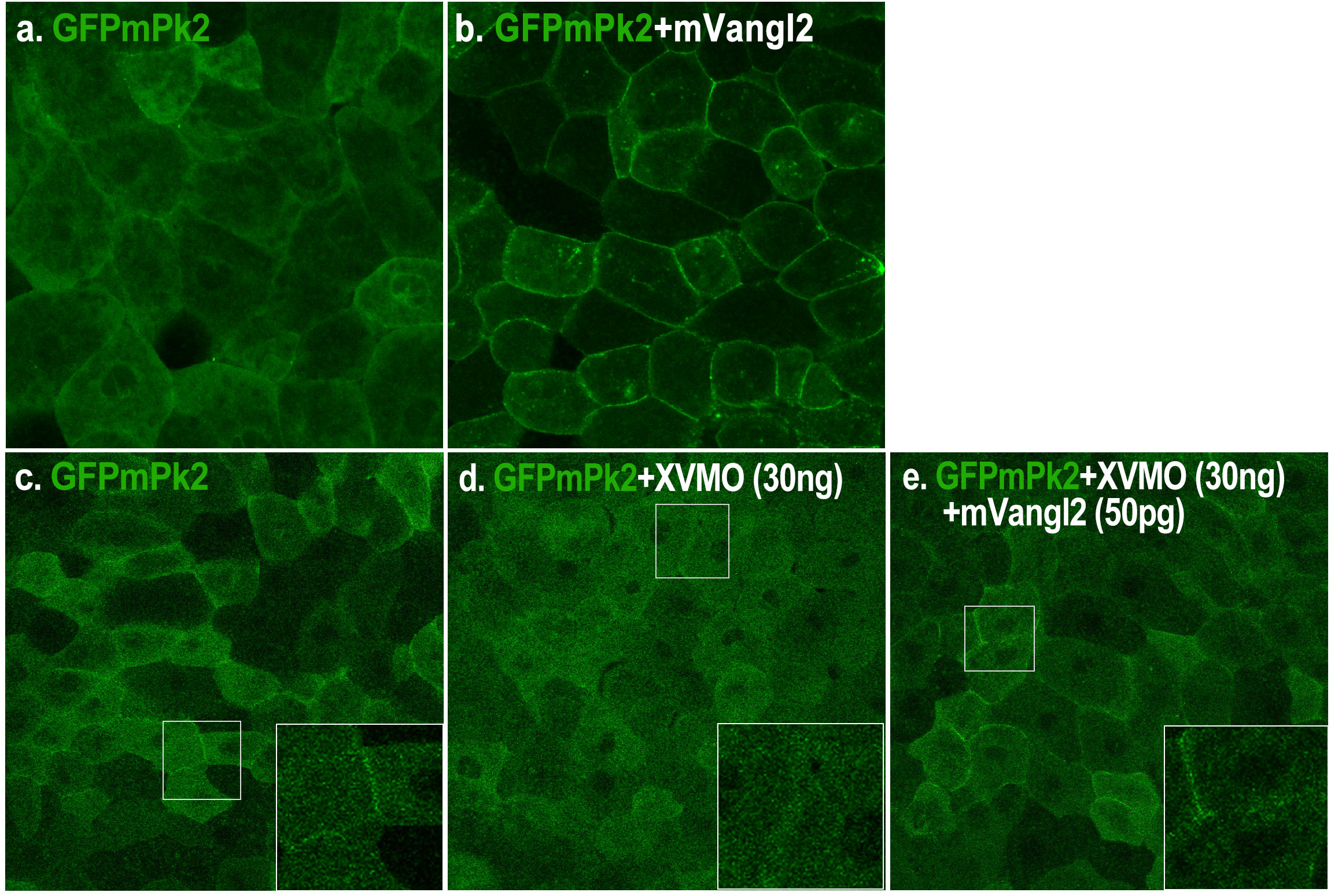
EGFP tagged mPk2 display uniform cytoplasmic distribution with weak enrichment at the plasma membrane when expressed in animal cap cells (a, c, and enlarged view in c). Plasma membrane enrichment of mPk2 is enhanced by co-injection of mVangl2 (b), but eliminated by morpholino knockdown of endogenous XVangl2 (d). Injecting a small amount of *mVangl2* (0.05 ng) could restore mPk2 plasma membrane enrichment in *XVangl2* morphant animal cap explants (e).

**Supplementary Figure 4.**
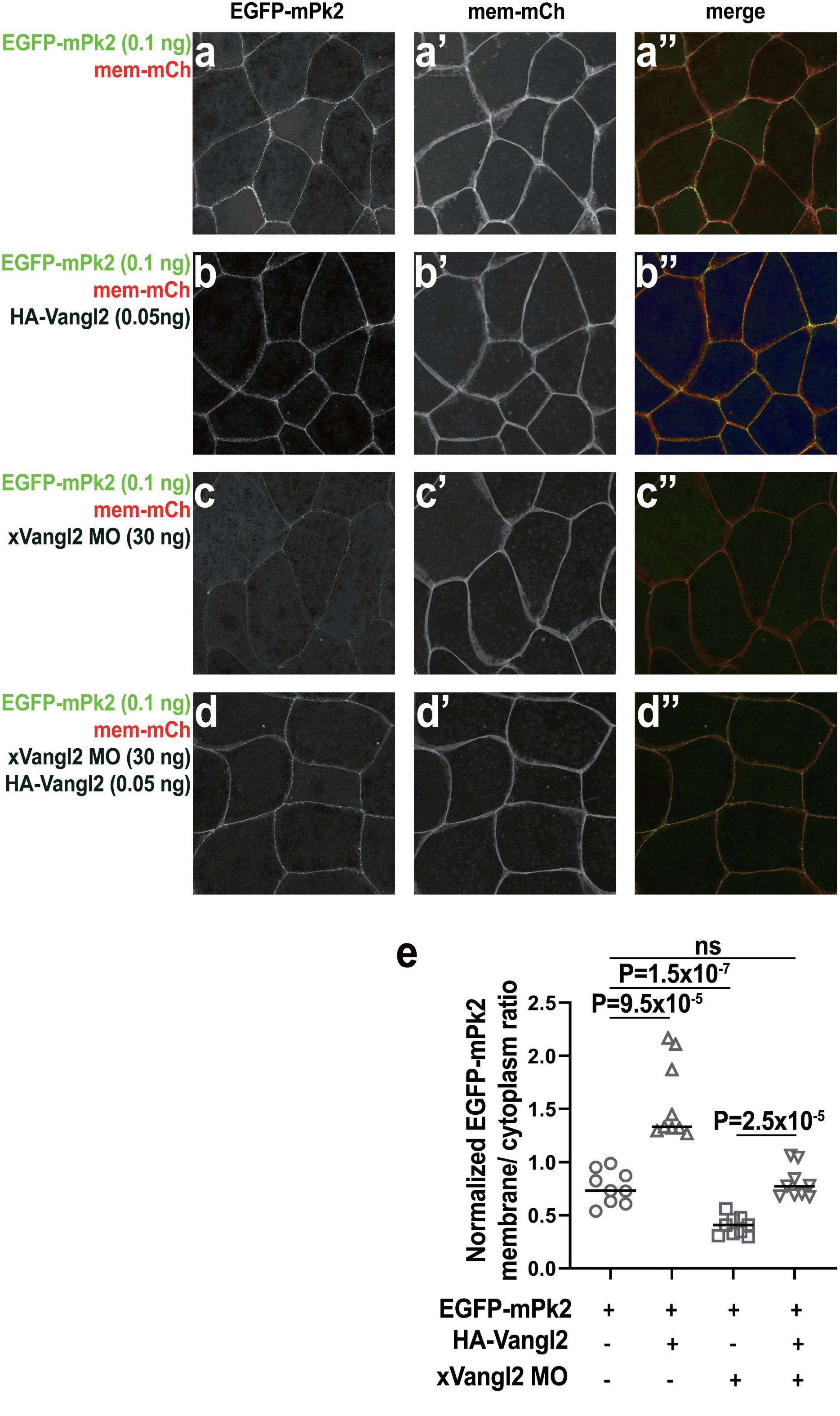
EGFP-mPk2 displayed both diffused cytoplasmic distribution and enrichment at the plasma membrane when injected into the DMZ (a-a”). Plasma membrane enrichment of mPk2 is enhanced by co-injection of 0.05 ng mVangl2 (b-b”), but diminished by knockdown of endogenous XVangl2 with 30 ng morpholino (c-c”). Co-injecting 0.05 ng *mVangl2* mRNA can rescue mPk2 plasma membrane enrichment in *XVangl2* morphants (d-d”). EGFP-mPk2 membrane enrichment was measured by calculating the “normalized EGFP-mPk2 membrane/ cytoplasm ratio” in (e). For this calculation, membrane mCherry (mem-mCh) was co-injected in each experiment to mark the plasma membrane, and mCh signal was used to segment the plasma membrane from the cytoplasm in ImageJ. The average signal intensities of EGFP-mPk2 and mem-mCh on the plasma membrane and in cytoplasm were then measured using ImageJ. The membrane-to-cytoplasm signal intensity ratio of EGFP-mPk2 is divided by that of mem-mCh to calculate the “normalized EGFP-mPk2 membrane/ cytoplasm ratio”. Each experiment was repeated with three separate injections on different days, and at least three embryos were injected each time for imaging analyses. Unpaired T-test was used to compare the “normalized mPk2 membrane/ cytoplasm ratio” of different groups, and the p vales are indicated between different groups.

**Supplementary Figure 5.**
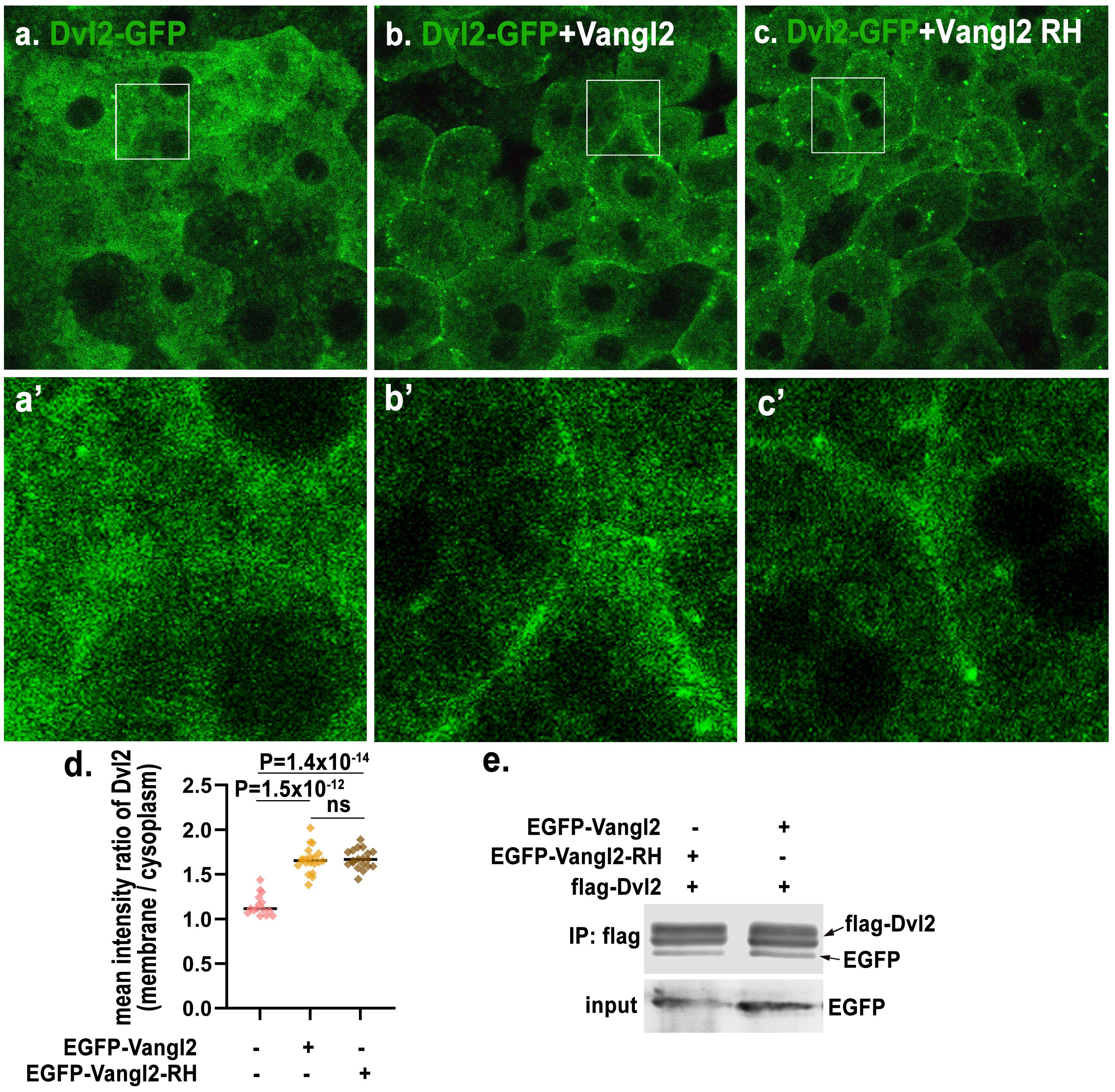
Dvl2-EGFP display uniform cytoplasmic distribution when expressed alone in *Xenopus* animal cap cells (a, a’), but co-expression with either wild-type Vangl2 (b, b’) or Vangl2 R177H variant (c, c’) recruits Dvl2-EGFP to the plasma membrane. (d) Quantification of the ratio of plasma membrane vs. cytoplasmic Dvl2-EGFP signal intensity in (a), (b) and (c). Co-IP and western blot did not reveal significant impact of R177H on either Vangl2 protein level or binding to Dvl2 (e).

**Supplementary Figure 6.**
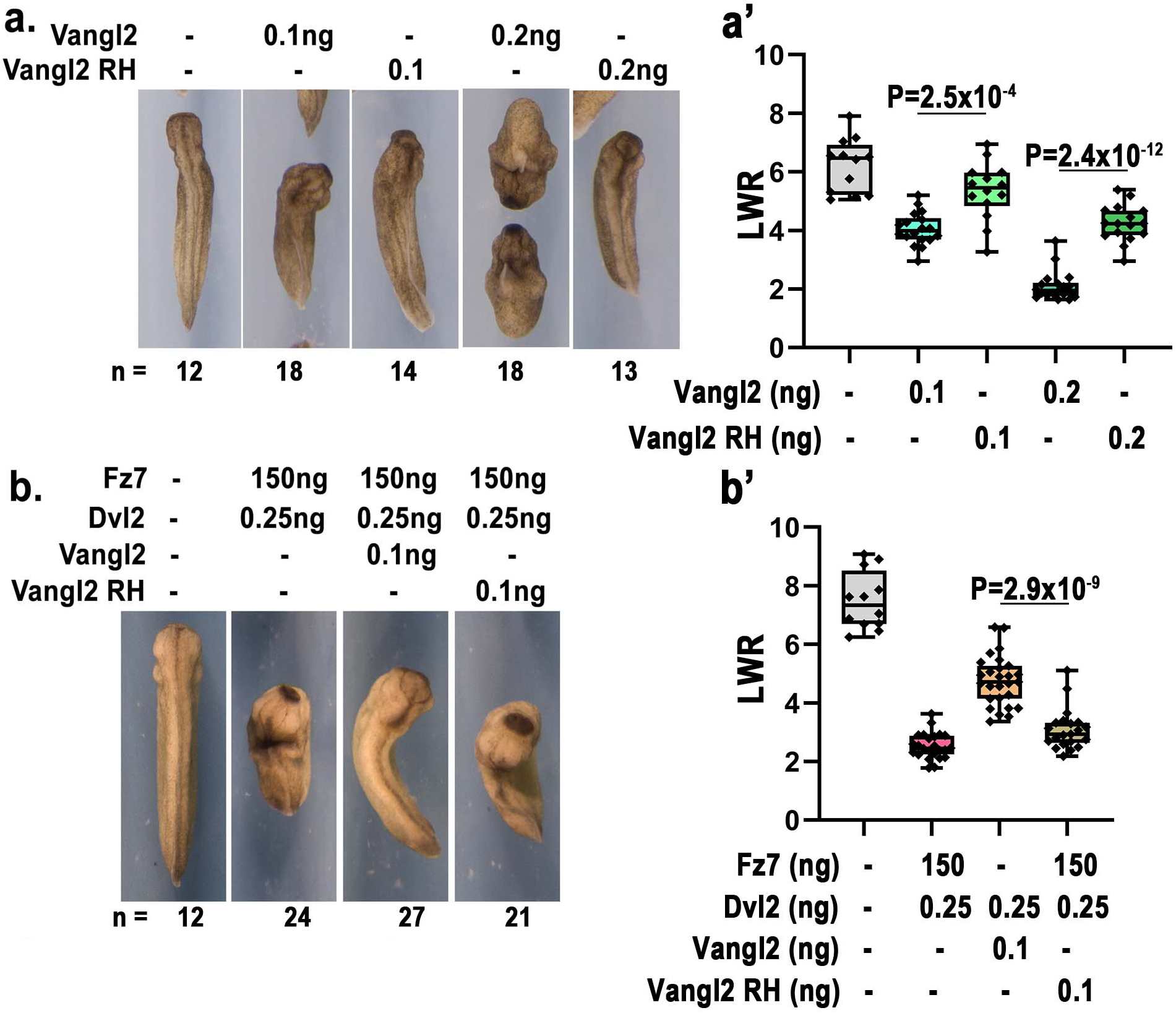
When injected into the DMZ, 0.2 ng and 0.1 ng wild-type mouse *Vangl2* mRNA causes severe and moderate CE defect, respectively. *Vangl2 R177H* mRNA, however, causes significantly milder CE defect (a). *Fz7* and *Dvl2* co-injection caused severe CE defects can be rescued by 0.1 wild-type *Vangl2*, but not by *Vangl2 R177H* (b). CE phenotype in (a) and (b) was determined by quantifying the length-to-width ratio (LWR) of the embryos in each group. Experiments were repeated three times and the total number of embryos analyzed is indicated below each panel in (a) and (b). Data are presented as box plots in (a’) and (b’), with the whiskers indicating the minima and maxima, the center lines representing median, the box upper and lower bounds representing 75th and 25th percentile, respectively. Two-tailed, unpaired T-test was used to compare the LWR of different groups, and the p vales are indicated in (a’) and (b’) between different groups.

**Supplementary Figure 7.**
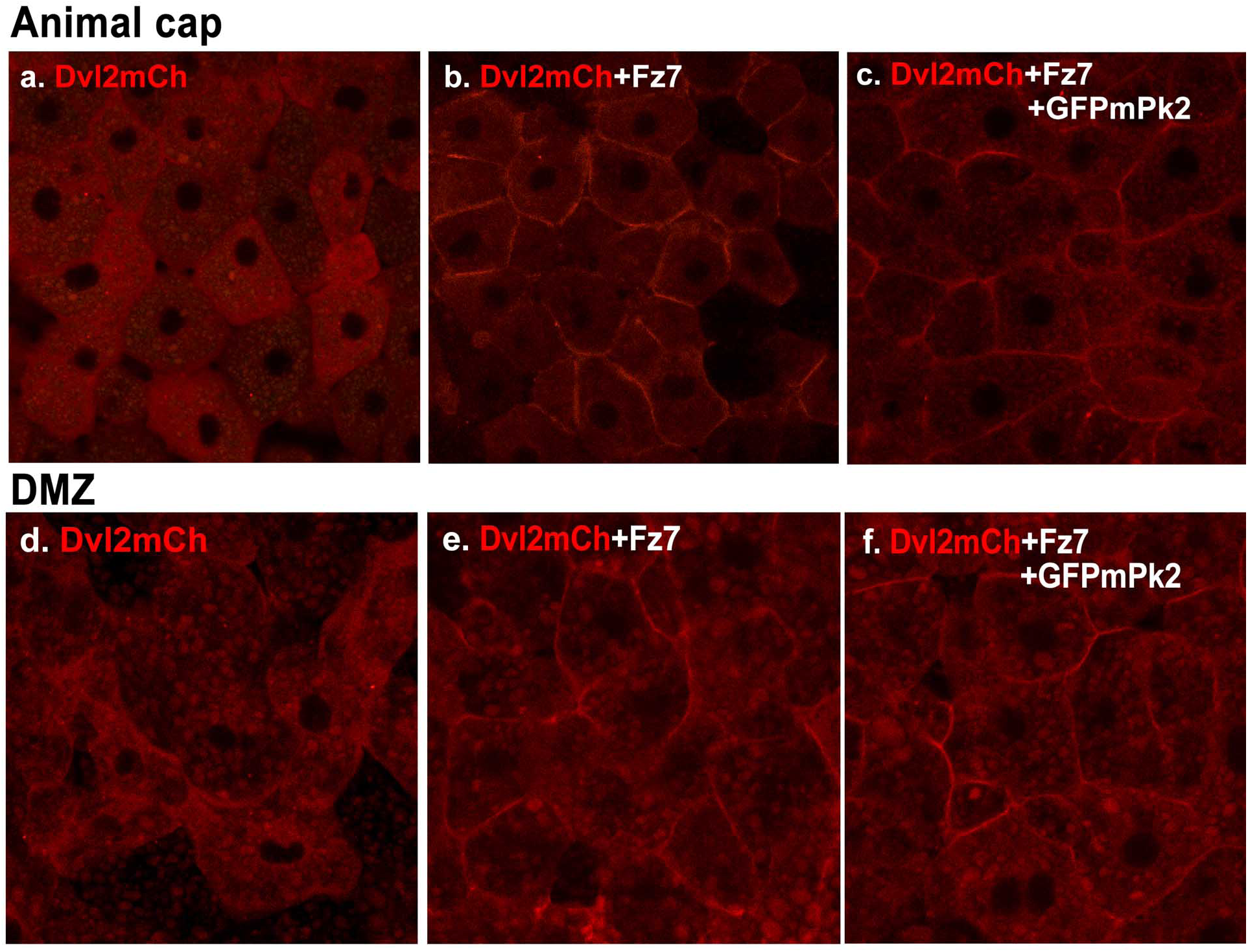
In both animal cap and DMZ cells, Fz7 can recruit Dvl2 from the cytoplasm to the plasma membrane (compare b to a, and e to d). Over-expression of Pk2 does not inhibit Fz7-mediated plasma membrane recruitment of Dvl2 (c, f).

**Supplementary Figure 8.**
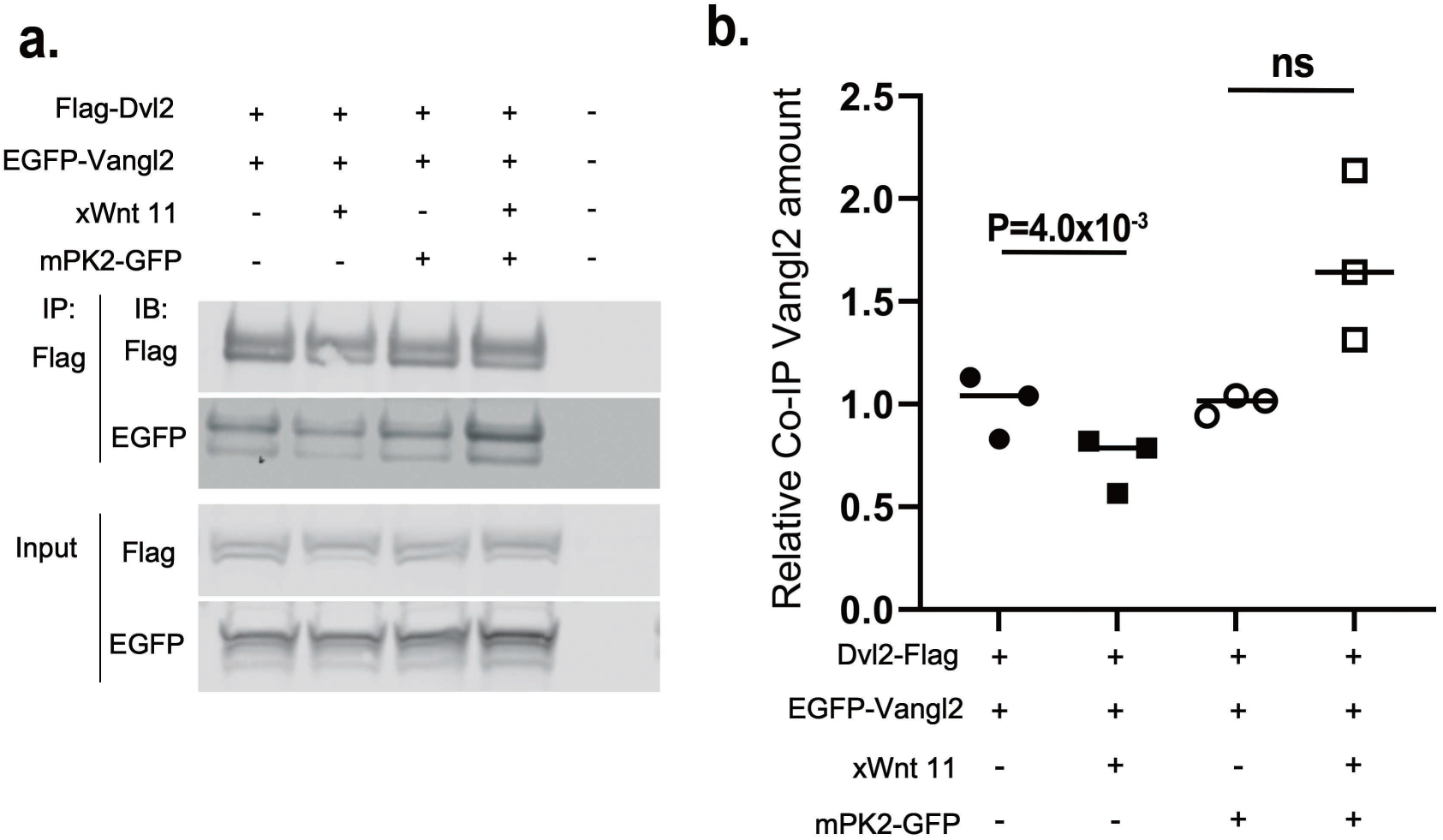
(a) In DMZ explant, binding between flag-Dvl2 and EGFP-Vangl2 is reduced by co-injection of Wnt11. But over-expression of Pk2 can prevent Wnt11-induced dissociation of Dvl2 and Vangl2. (b) Quantification of Vangl2 co-IP over pulled down Flag-Dvl2 from 3 separate experiments.

**Supplementary Figure 9.**
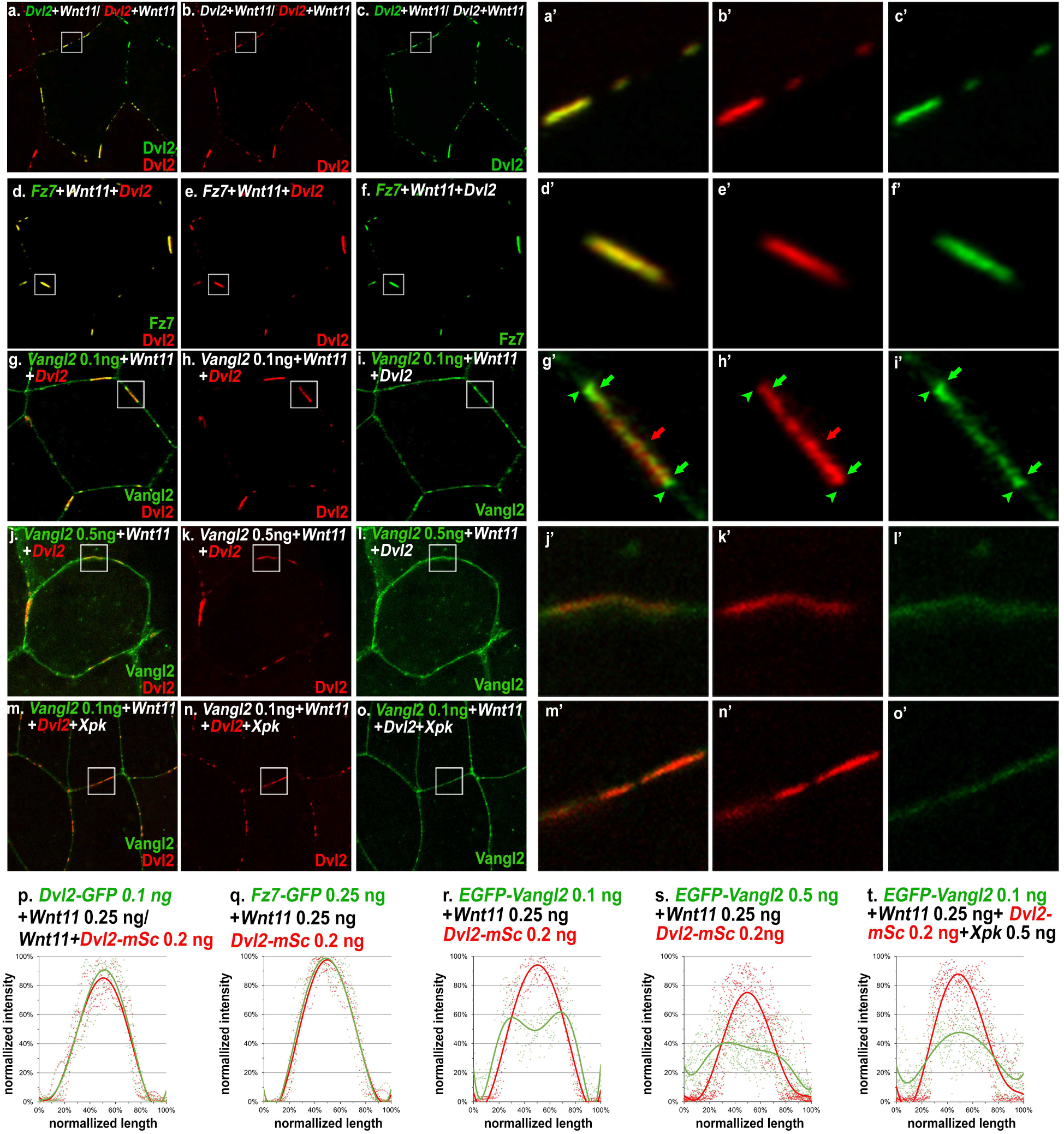
In DMZ explants, 0.1 ng of EGFP-tagged mouse *Dvl2* (*Dvl2-EGFP*) and 0.2 ng of mScarletI-tagged mouse *Dvl2* (*Dvl2-mSc*) are separately injected into two adjacent blastomeres with *Wnt11.* Wnt11-induced Dvl2-mSc and Dvl2-EGFP patches are both observed, and they are aligned along the border of adjacent cells (a-c, and enlarged view of the boxed area in a’-c’). Co-injection of *Dvl2-mSc* with *Fz7-EGFP* indicates complete overlap between Dvl2 and Fz7 in the patches (d-f’). In contrast, Vangl2, when expressed at moderate levels (0.1 ng), is distributed more broadly along the plasma membrane (g-i), but also displays enrichment immediately outside and at the edge of Dvl2 patches (g’-I’, arrowheads and arrows, respectively). High level of *Vangl2* injection (0.5 ng) perturbs Wnt11-induced Dvl2 patch formation and makes Dvl2 more evenly distributed with Vangl2 (j-l’). The same effect can also be achieved by co-expressing Pk with moderate level of Vangl2 (m-o’). (p-t) Measurement of the relative intensity of Dvl2-mSc along the patches with either Dvl2-EGFP, Fz7, Vangl2 at moderate (0.1 ng) and high (0.5 ng) levels, or Vangl2 (0.1 ng) and XPk (0.5 ng) co-injection.

**Supplementary Figure 10.**
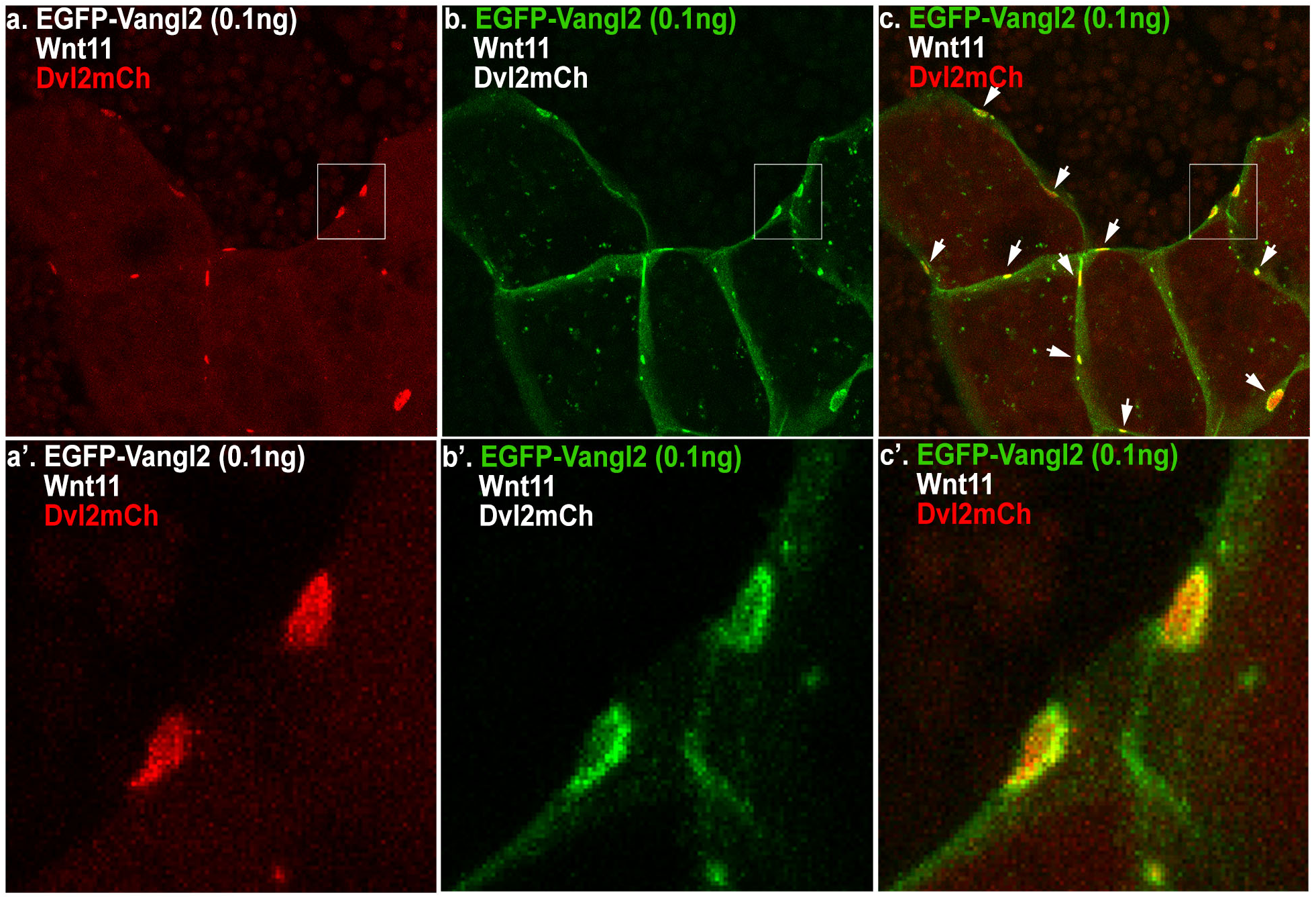
Moderate level of *EGFP-Vangl2* (0.1 ng) was co-injected with *Wnt11* and mCherry tagged *Dvl2*. 3D reconstruction of confocal images shows that under this condition, all Dvl2 proteins on the plasma membrane formed discrete patches at the apical cell-cell junctions (a), while Vangl2 was distributed more diffusely on the plasma membrane (b) with enrichment overlapping with Dvl2 patches (c, white arrows). Enlarged views (a’-c’) show that enriched Vangl2 forms rings that encircle Dvl2 patches.

**Supplementary Figure 11.**
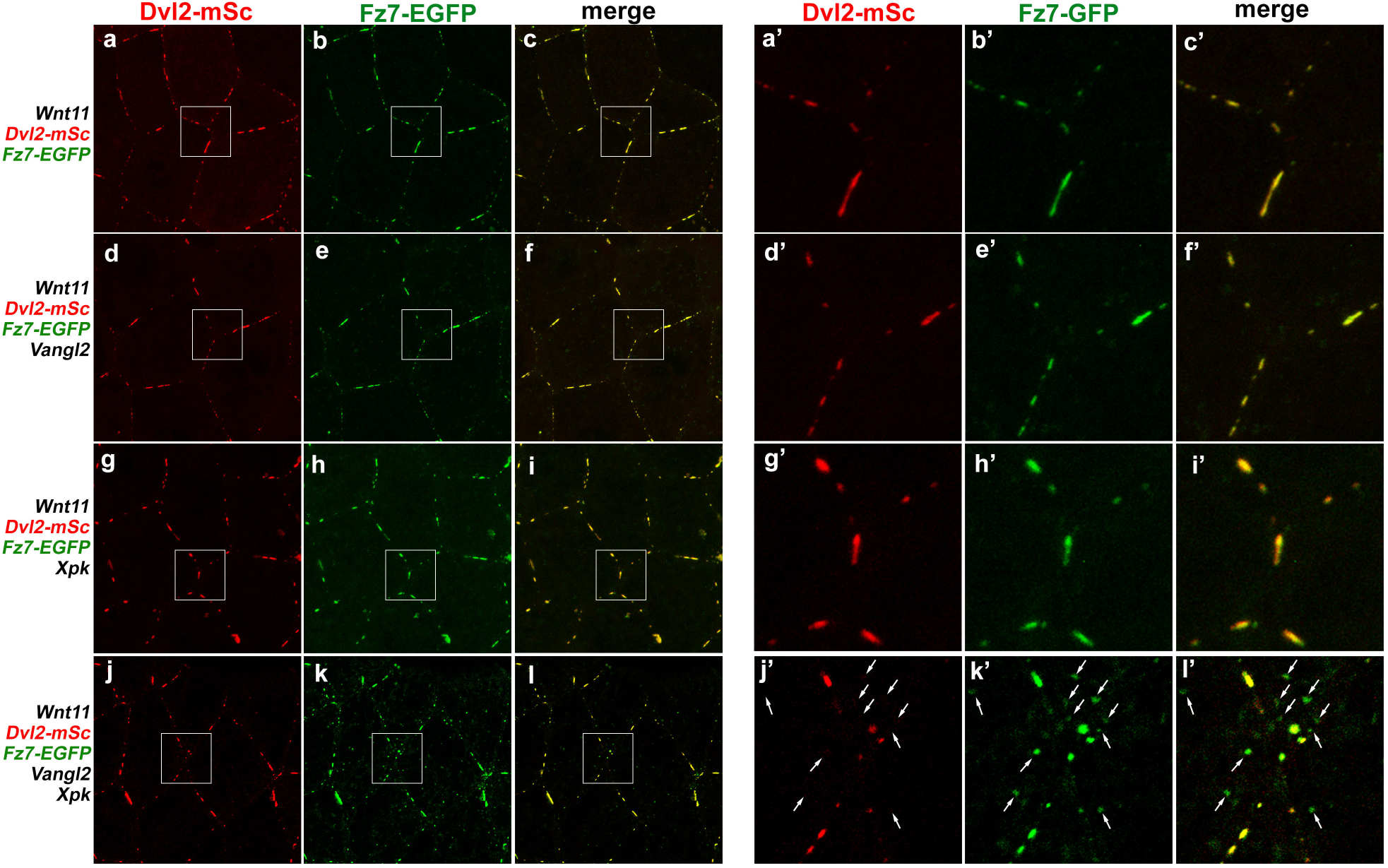
In DMZ explants, Wnt11 induces formation of overlapping Fz7-EGFP and Dvl2-mScarletI (Dvl2-mSc) patches at the cell-cell contact (a-c’). These patches are not affected by co-injecting moderate amount of Vangl2 (d-f’, 0.1 ng mRNA) or XPk (g-i’, 0.5 ng mRNA) individually. Vangl2 and Pk co-expression, however, not only diminished Dvl2 patches but also dispersed Fz7 into small puncta (j-l). Enlarged views revealed that some of the Fz7 puncta are on the plasma membrane and remain co-localized with Dvl2, while the others are located in the cytoplasm near the plasma membrane (k’’, l’, arrows) and contain only Fz7 but not Dvl2 (compare arrows in j’ to k’).

**Supplementary Figure 12.**
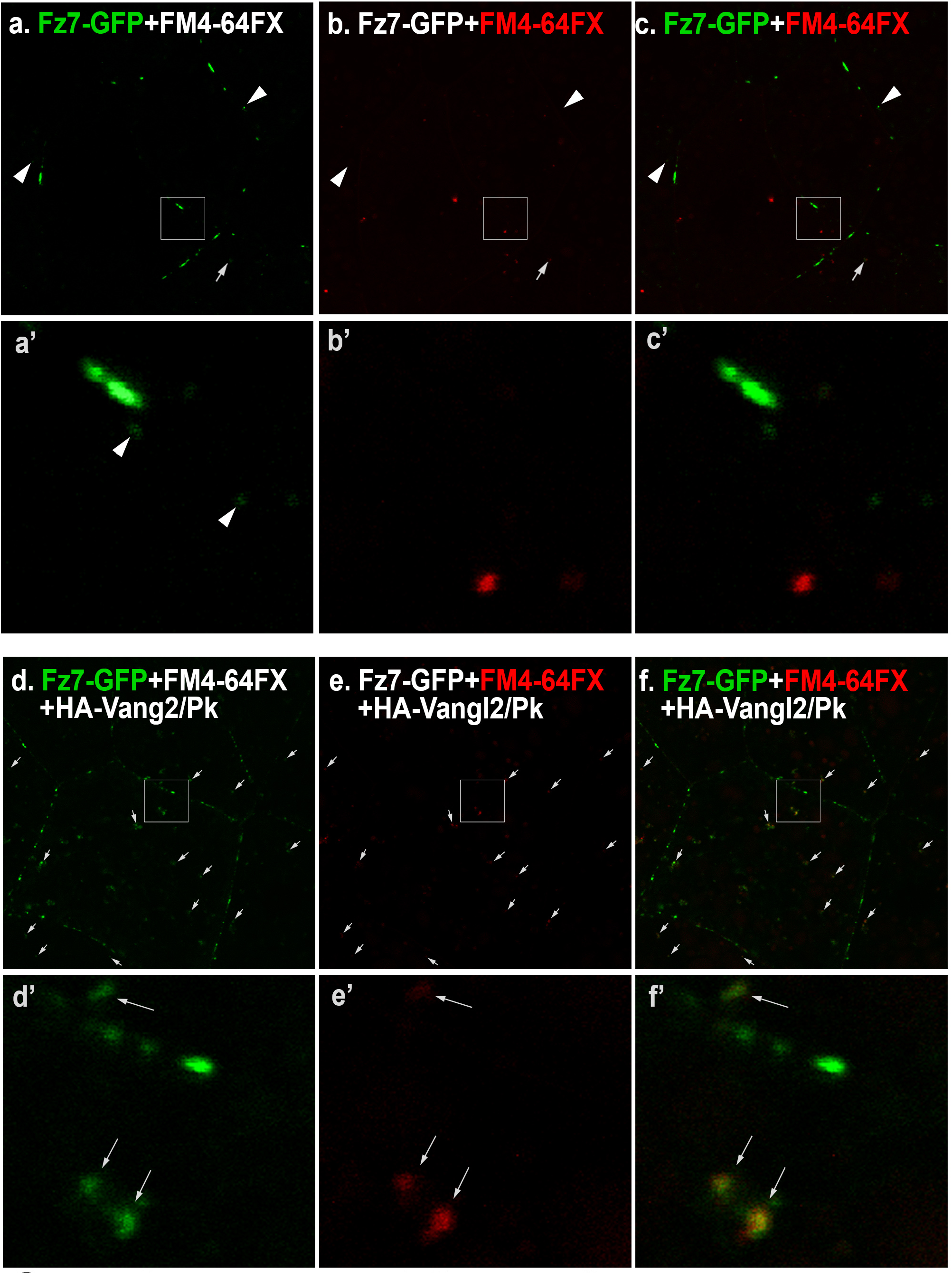
*Xenopus* explants were incubated with FM4-64FX, a fluorescent dye that is membrane impermeable and can only be internalized through endocytosis. In explants in which *Fz7-GFP* was co-injected only with *Wnt11*, Fz7 primarily formed patches on the cell cortex. Most of the internalized FM4-64FX puncta were devoid of Fz7, and the few Fz7 puncta were devoid of FM4-64FX (arrowheads in a-c, and enlarged views a’-c’). In contrast, in explants in which *Fz7-GFP* and *Wnt11* were co-injected with *Vangl2/Pk*, majority of the Fz7-GFP puncta are positive for FM4-64FX (arrows in d-f and enlarged views d’-f’), indicating that they were largely formed through endocytosis.

**Supplementary Figure 13.**
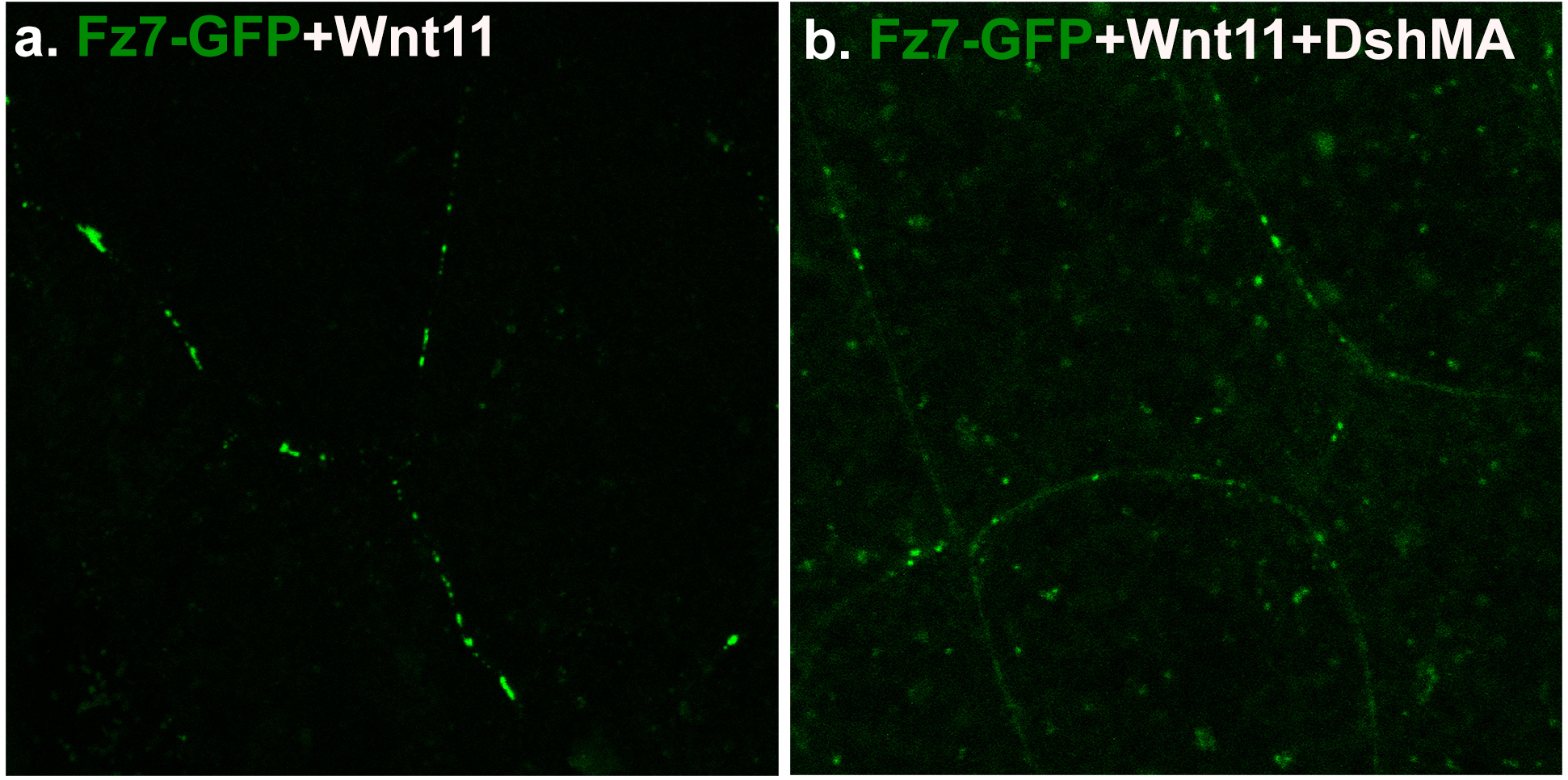
Wnt11 induces formation of Fz7 patches along the plasma membrane (a). Over-expression of DshMA, a mitochondria tethered form of Dvl that can sequester endogenous Dvl/Dsh away from the plasma membrane, inhibits Fz7 patch formation, reduces membrane level of Fz7, and results in Fz7 cytoplasmic puncta (b).

**Supplementary Figure 14.**
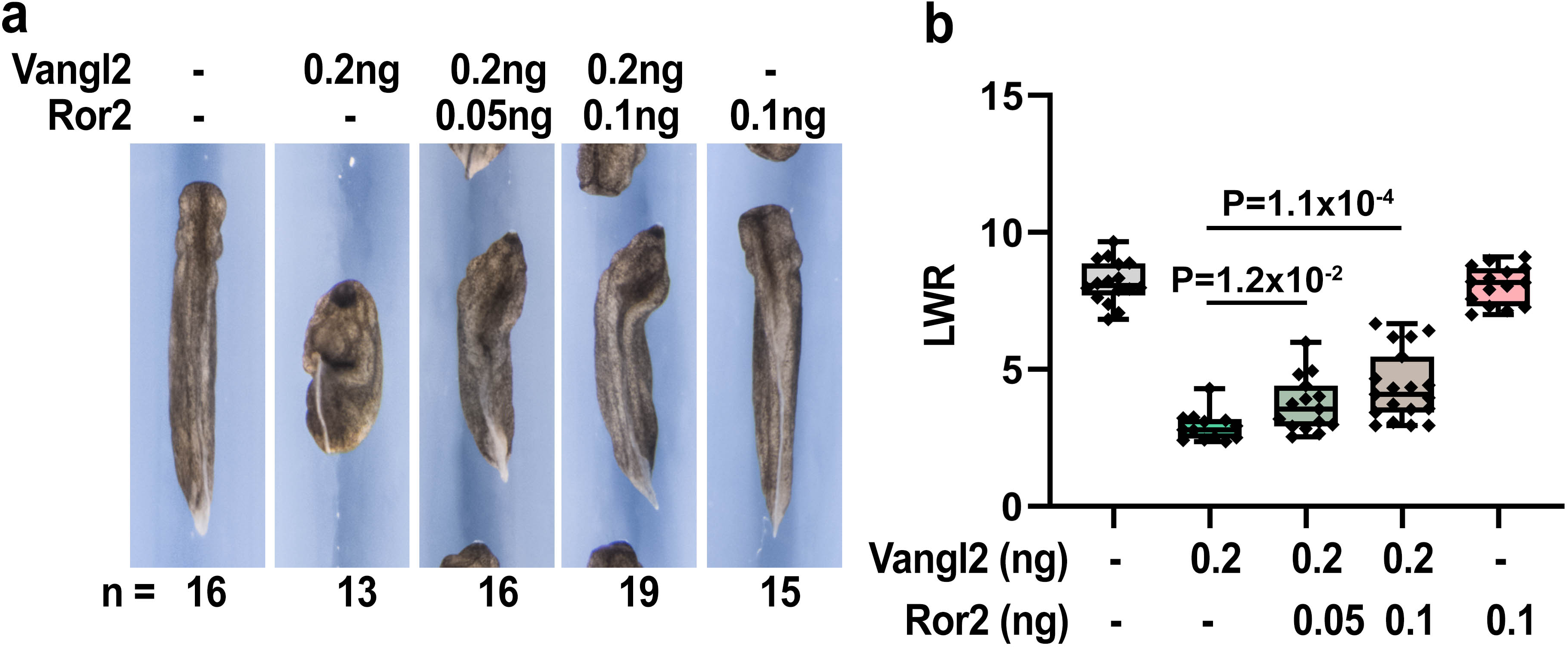
DMZ injection of 0.2 ng of *Vangl2* mRNA induces severe CE defects, which can be rescued by co-injecting 0.05 or 0.1 ng of *Xror2* mRNA (a). CE phenotype in (a) was determined by quantifying the length-to-width ratio (LWR) of the embryos in each group. Experiments were repeated three times and the total number of embryos analyzed is indicated below each panel in (a). Data are presented as box plots in (b), with the whiskers indicating the minima and maxima, the center lines representing median, the box upper and lower bounds representing 75th and 25th percentile, respectively. Two-tailed, unpaired T-test was used to compare the LWR of different groups, and the p vales are indicated in (b) between different groups.

**Supplementary Figure 15.**
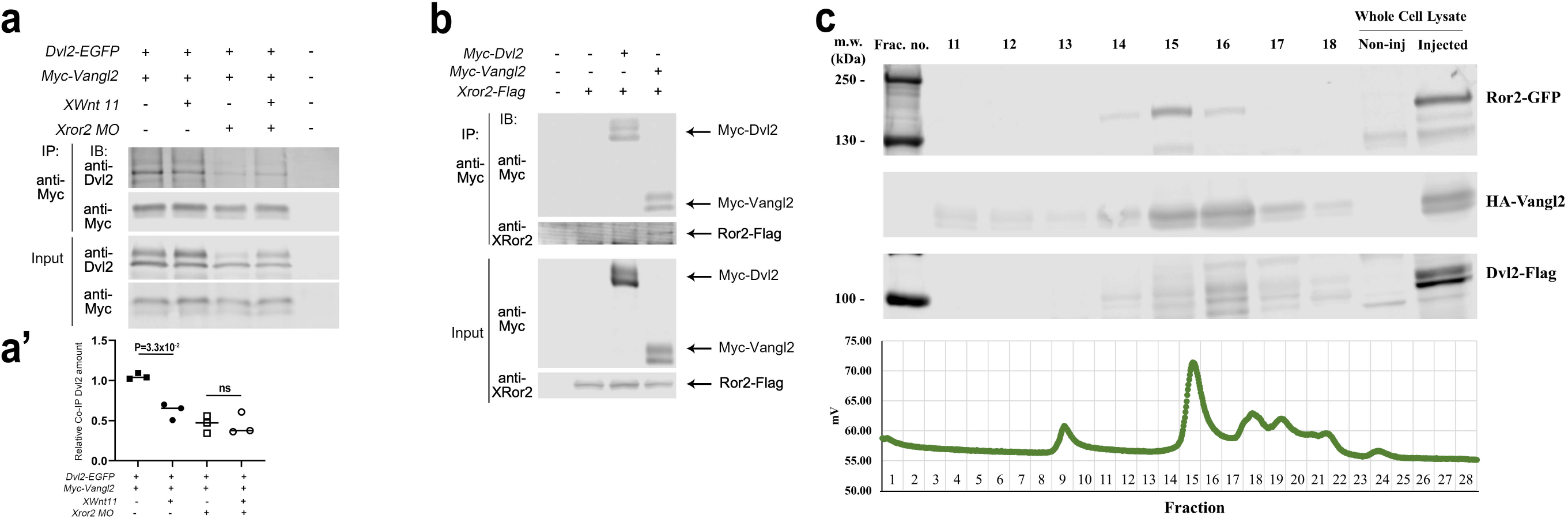
In *Xenopus* embryo extract, co-IP experiment shows that binding between Dvl2-EGFP and Myc-Vangl2 is reduced by Wnt11 (compare lane2 and 3). Knocking down endogenous XRor2 with morpholino (*Xror2 MO*) can prevent Wnt11-induced dissociation of Dvl2 and Vangl2 (compare lane 4 and 5) (a). (a’) Quantification of Dvl2-EGFP co-IP against Myc-Vangl2 pull down (n=3). Note that Dvl2-Vangl interaction is reduced with XRor2 knock down, either with or without Wnt11 co-injection. (b) Co-IP experiment shows that Ror2-Flag can be pulled down by co-injected Myc-Vangl2 but not Myc-Dvl2. (c) Fluorescence-detection size exclusion chromatography (FSEC) with protein extract from *Xenopus* embryos injected with *Xror2-EGFP*, *HA-Vangl2* and *Flag-Dvl2*. A peak of EGFP signal is detected around fraction 15 (lower panel). Western blot analyses show co-fractionation of Ror2-EGFP, HA-Vangl2 and Flag-Dvl2 in fractions 14 (approximate molecular weight 773-1717 kD), 15 (∼348-773 kD) and 16 (∼166-348 kD).

**Supplementary Figure 16.**
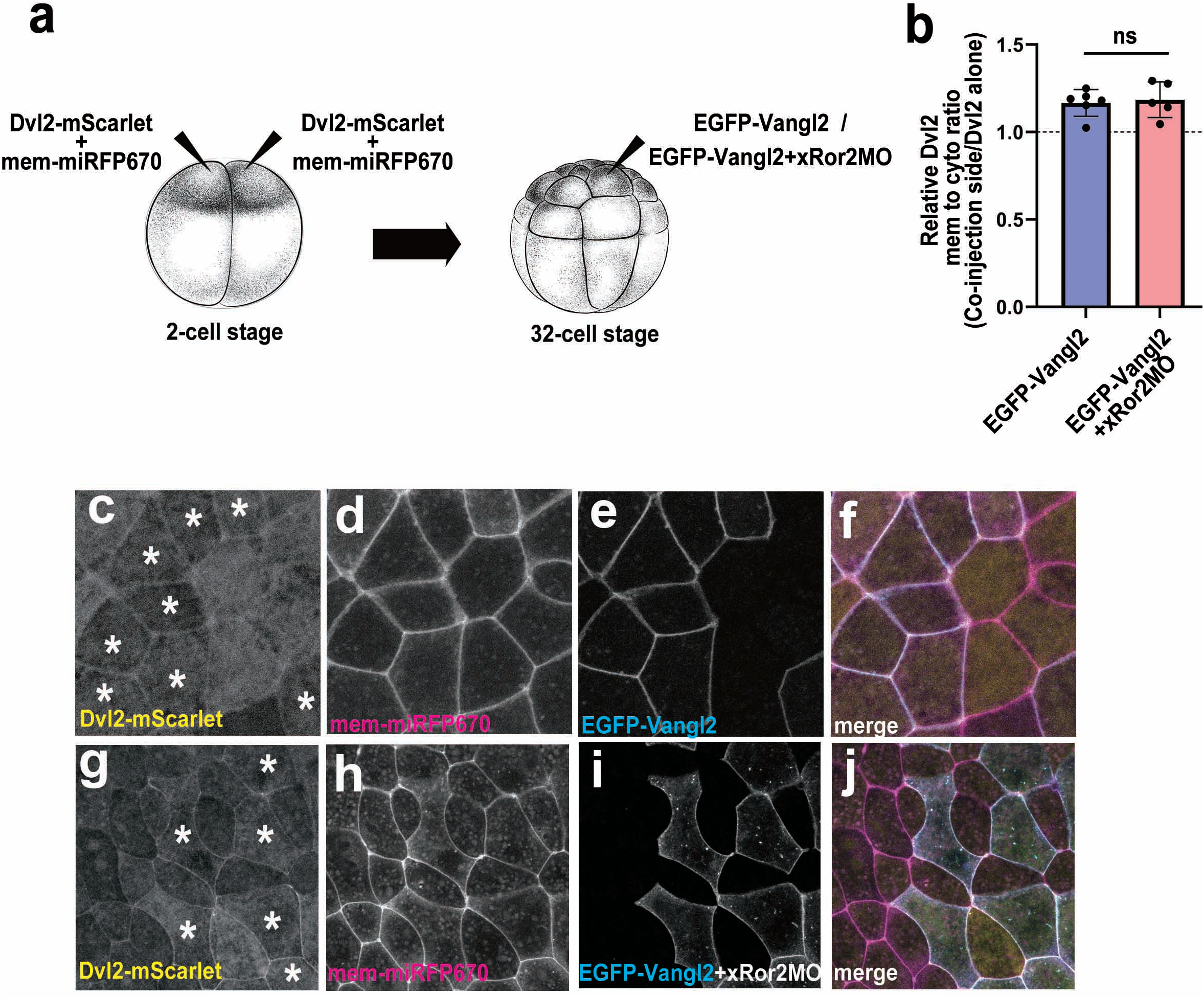
(a) Scheme of making chimeric embryos to examine Dvl2 plasma membrane recruitment by Vangl2 with or without Ror2 knockdown. 2-cell stage embryos receive the first injection into the animal side of both blastomeres, with mRNA encoding Dvl2-mScarlet-I and mem-miRFP670. The embryos are cultured to 32-cell stage, and receive a second injection into a single animal pole blastomere with mRNA encoding EGFP-Vangl2, or EGFP-Vangl2 with Ror2 MO. The embryos are cultured to stage 8, and the animal caps are dissected for imaging. (b) Dvl2 plasma membrane recruitment by Vangl2 is measured by the average plasma membrane-to-cytoplasm intensity ratio of Dvl2 in cells co-expressing Dvl2 and Vangl2, normalized by that in cells expressing Dvl2 alone. (c-f) Image of animal cap explant from embryos injected with EGFP-Vangl2 at 32-cell stage. (g-j) Image of animal cap explant from embryos injected with EGFP-Vangl2 and Ror2 MO at 32-cell stage.

**Supplementary Figure 17.**
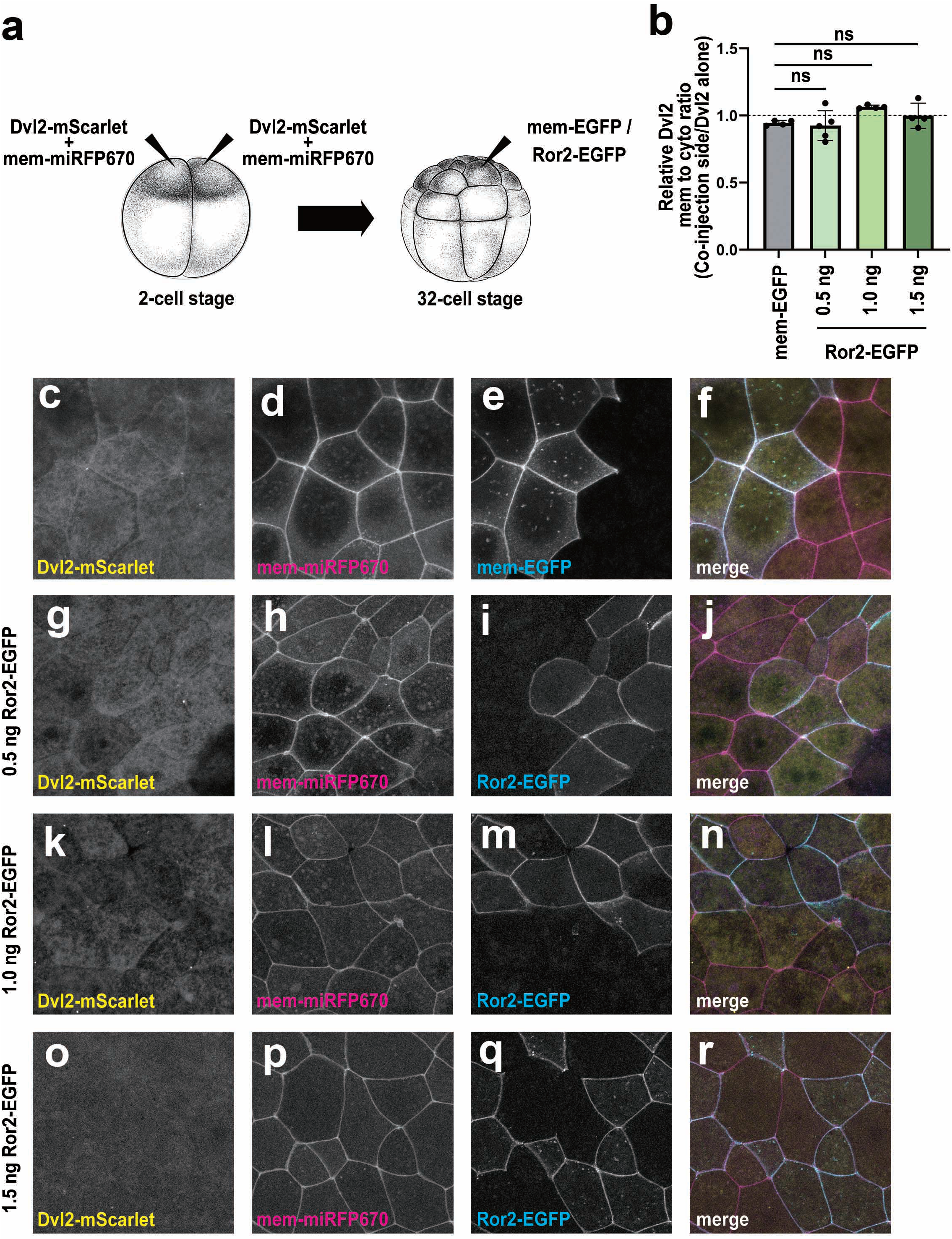
(a) Scheme of making chimeric embryos to examine Dvl2 plasma membrane recruitment by Ror2. 2-cell stage embryos receive the first injection into the animal side of both blastomeres, with mRNA encoding Dvl2-mScarlet-I and mem-miRFP670, a plasma membrane marker. The embryos are cultured to 32-cell stage, and receive a second injection into a single animal pole blastomere with mRNA encoding either mem-EGFP or varying levle of Ror2-EGFP. The embryos are cultured to stage 8, and the animal caps are dissected for imaging. (b) Dvl2 plasma membrane localization is measured by the average plasma membrane-to-cytoplasm intensity ratio of Dvl2 in cells co-expressing Dvl2 and Ror2 (or mem-EGFP control), normalized by that in cells expressing Dvl2 alone. (c-f) Image of animal cap explant from embryos injected with mem-EGFP control at 32-cell stage. (g-r) Image of animal cap explant from embryos injected with 0.5 ng (g-j), 1 ng (k-n) or 1.5 ng (o-r) of Ror2-EGFP mRNA at 32-cell stage.

**Supplementary Figure 18.**
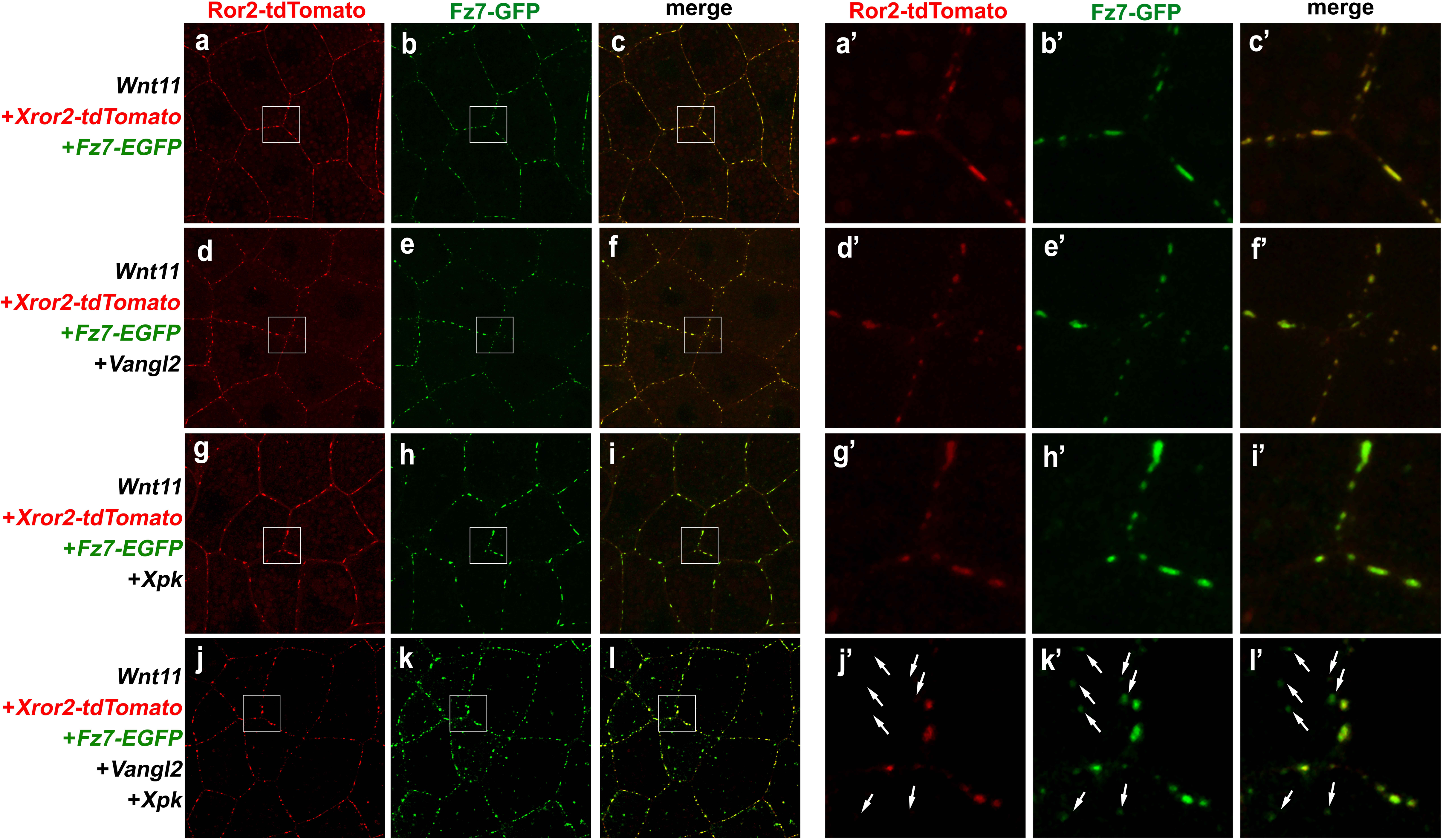
(a-c) In DMZ explants, Wnt11 induces co-injected XRor2-tdTomanto and Fz7-EGFP to co-cluster into patches on the cell cortex. Overexpression of moderate level of Vangl2 (d-f, 0.1 ng) or Pk (g-I, 0.5 ng) individually does not perturb co-clustering of Ror2 with Fz7 into patches in response to Wnt11, but their co-overexpression (j-l) diminishes Ror2-Fz7 patches into puncta and cause Fz7 to form cytoplasmic puncta whereas Ror2 remained on the plasma membrane (compare arrows in j’-l’). a’-l’ are enlarged view of a-l, respectively.

## Notes

### Competing Interest Statement

The authors have declared no competing interest.

### Summary of Updates

Addition of Fig. 1h, h'; Addition of Suppl. Fig. 16 and 17; Minor revision to the manuscript

